# Targeting an evolutionarily conserved “E-L-L” motif in the spike protein to develop a small molecule fusion inhibitor against SARS-CoV-2

**DOI:** 10.1101/2022.03.16.484554

**Authors:** Indrani Das Jana, Prabuddha Bhattacharya, Karthick Mayilsamy, Saptarshi Banerjee, Gourab Bhattacharje, Sayan Das, Seemanti Aditya, Anandita Ghosh, Andrew R. McGill, Syamanthak Srikrishnan, Amit Kumar Das, Amit Basak, Shyam S. Mohapatra, Bala Chandran, Devesh Bhimsaria, Subhra Mohapatra, Arunava Roy, Arindam Mondal

## Abstract

As newer variants of SARS-CoV-2 continue to pose major threats to global human health and economy, identifying novel druggable antiviral targets is the key towards sustenance. Here, we identify an evolutionary conserved “E-L-L” motif present within the HR2 domain of all human and non-human coronavirus spike (S) proteins that play a crucial role in stabilizing the post-fusion six-helix bundle (6-HB) structure and thus, fusion-mediated viral entry. Mutations within this motif reduce the fusogenicity of the S protein without affecting its stability or membrane localization. We found that posaconazole, an FDA-approved drug, binds to this “E-L-L” motif resulting in effective inhibition of SARS-CoV-2 infection in cells. While posaconazole exhibits high efficacy towards blocking S protein-mediated viral entry, mutations within the “E-L-L” motif rendered the protein completely resistant to the drug, establishing its specificity towards this motif. Our data demonstrate that posaconazole restricts early stages of infection through specific inhibition of membrane fusion and viral genome release into the host cell and is equally effective towards all major variants of concerns of SARS-CoV-2 including beta, kappa, delta, and omicron. Together, we show that this conserved essential “E-L-L” motif is an ideal target for the development of prophylactic and therapeutic interventions against SARS-CoV-2.

## Introduction

The COVID-19 pandemic, caused by the Severe Acute Respiratory Syndrome Coronavirus 2 (SARS-CoV-2), has exposed the global unpreparedness in controlling infectious outbreaks caused by coronaviruses (https://covid19.who.int/, 2022). Though vaccines and repurposed drugs are being administered through emergency use authorization (Mcgill et al., 2021), the world is still suffering from consecutive waves of the pandemic caused by newer variants of SARS-CoV-2, generated through acquired mutations in the spike and other viral proteins (https://www.who.int/en/activities/tracking-SARS-CoV-2-variants/, 2022). To outcompete this rapidly evolving pathogen, it is crucial to develop new intervention strategies by focusing on highly conserved molecular targets present within the virus, which will target both the current as well as any forthcoming variants.

In viruses of the *Coronaviridae* family, the spike protein is responsible for receptor recognition and membrane fusion leading to the release of the viral genomic RNA into the host cell, which makes it a prime target for the development of novel antiviral drugs (Hofmann & Pöhlmann, 2004; Letko et al., 2020). The 1273 aa SARS-CoV-2 S is proteolytically cleaved into two different subunits, S1 and S2 (Figure 1A), where the S1 subunit consists of an N-terminal and a receptor-binding domain (RBD) which is responsible for binding to the cell surface ACE2 receptors. The S2 subunit, consists of the fusion peptide (FP), heptapeptide repeat sequences 1 and 2 (HR1 and HR2), transmembrane domain (TM), and the cytoplasmic domain (CD) which together drives membrane fusion (Y. Huang et al., 2020). On the virion surface, the S protein exists as a homotrimer where the S1 and S2 fragments remain non-covalently attached in a pre-fusion metastable conformation (Wrapp et al., 2020). Upon host cell receptor binding, the S2 subunit undergoes major structural rearrangements from the pre-fusion to a post-fusion conformation, which results in protrusion of the hydrophobic FP into the host cell membrane, close juxtaposition of the viral and host cell membranes and finally fusion of the two membranes together (Bosch et al., 2003).

**Figure 1.**
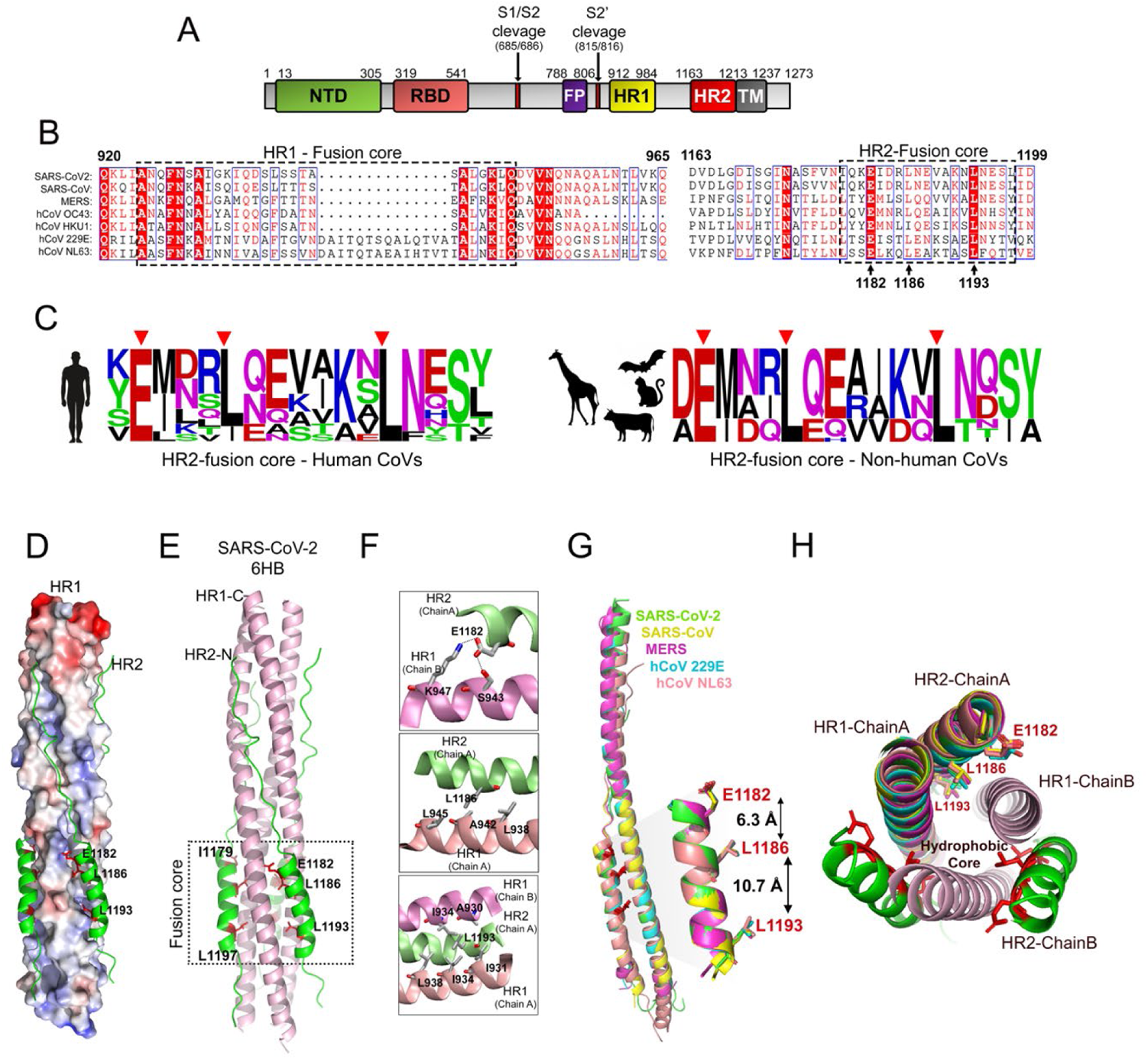
An evolutionary conserved “E-L-L” motif in the HR2 domain of SARS-CoV-2 S: (A) Domain organization of SARS-CoV-2 Spike protein. (B) Multiple sequence alignment of HR1 and HR2 domains of spike proteins from different human infecting coronaviruses with the fusion-core region highlighted by dashed rectangles. Amino acid residue numbers correspond to the SARS-CoV-2 S protein. (C) Logo plot representing amino acid frequencies in the fusion-core region of the HR2 from human or different animal infecting coronaviruses, highlighting the conserved E1182, L1186, and L1193 residues. (D) SARS-CoV-2 spike protein HR1 (surface charge representation) and HR2 (in ribbon representation) in the form of six-helix bundle (PDB ID: 6LXT). (E) Ribbon representation of three HR1 (light pink) and three HR2 (green) forming the 6-HB bracketed by the dashed rectangle. The E1182, L1186, and L1193 residues are highlighted in red stick representations. (F) Top. E1182 of HR2 from chain A (lime green) forming a salt bridge interaction and hydrogen bond with K947 and S943 respectively of the HR1 of chain B (pink). Middle. Intramolecular hydrophobic interactions between L1186 of HR2 (chain A) and L-945, A-942, and L938 of HR1 (chain A, pale pink). Bottom. Intramolecular hydrophobic interactions between L1193 of HR2 Chain A and L938, I934 and I931 of HR1 chain A (pale pink), and A930 and I934 of HR1 chain B (pink). (G) Structural alignment of one pair of HR1-HR2 helices of different human infecting coronaviruses-green, SARS-CoV-2 (6LXT), yellow, SARS-CoV (1WYY), magenta, MERS-CoV (4NJL), cyan, 229E (5YL9), and, pale orange, NL63 (2IEQ). The zoomed image represents the aligned HR2 fusion-core regions along with the side chains of the E1182, L1186, and L1193 residues. (H) Top view of the 6-HB structure of SARS-CoV-2 spike with one pair of the helix structurally aligned with that of the other coronaviruses as mentioned in G.

In the S trimer, the HR1 and HR2 helices from three different S2 subunits interact with each other to form a rod-shaped six-helix bundle (6-HB), also known as “fusion-core”. In the 6-HB, three parallel HR1 helices from three individual protomers form a trimeric coiled-coil hydrophobic core around which the HR2s fold back and intertwine in an antiparallel manner (Shuai Xia, Liu, et al., 2020) participating in a series of intra- as well as intermolecular interactions. These interactions are critical for the stability of the post-fusion conformation and are therefore vital for the membrane fusion process (Y. Huang et al., 2020). This is why peptides that competitively inhibit HR1-HR2 interactions can effectively block 6-HB formation and hence, viral fusion (Lan et al., 2021; Outlaw et al., 2020; S Xia et al., 2019; Shuai Xia, Liu, et al., 2020; Shuai Xia, Zhu, et al., 2020; Y Zhu et al., 2020; Yuanmei Zhu et al., 2021). Although these studies highlight the importance of the non-covalent interactions in the fusion process, the use of peptides as an effective therapeutic drug has several limitations compared to small molecule inhibitors, which are often more potent, stable, have better pharmacokinetics, and are easier to produce (Fosgerau & Hoffmann, 2015; Otvos & Wade, 2014). However, to date, the development of small molecule fusion inhibitors against HCoVs have been remarkably limited, pointing towards the urgency of developing such molecules to treat SARS-CoV-2 and other related coronavirus infections.

Here, we used a rational drug repurposing approach to identify an FDA-approved small molecule that has the potential to interfere with the HR1-HR2 interaction and hence act as a broad-spectrum fusion inhibitor against different SARS-CoV-2 variants that are circulating worldwide. First, we identified a novel, evolutionary conserved “Ex_3_Lx_6_L” motif (abbreviated as E-L-L) in the HR2 domain of SARS-CoV-2 S and demonstrated that it plays a critical role in 6-HB formation and in fusion mediated viral entry, thereby establishing it as a potential target for fusion inhibitors. Subsequent *in-silico* screening led to the identification of posaconazole, a triazole antifungal drug (Schlossberg & Samuel, 2017), which we show, interacts with the “E-L-L” motif with high affinity. Posaconazole efficiently blocked membrane fusion, inhibiting SARS-CoV-2 S pseudovirus entry in host cells. Interestingly, pseudoviruses harboring mutations within the “E-L-L” motif of the S protein were resistant towards posaconazole, confirming (i) importance of the “E-L-L” motif in viral entry and (ii) high specificity of the drug towards the “E-L-L” motif mediated membrane fusion. Finally, we demonstrated the efficacy of posaconazole in inhibiting SARS-CoV-2 infection in Caco-2 cells through specific inhibition of membrane fusion, syncytium formation, and entry of viral genomic RNA into the host cells. Posaconazole also showed undiminished activity against the highly contagious beta (B.1.351), kappa (B.1.617.1) delta (B.1.617.2) and omicron (B.1.1.529) variants of SARS-CoV-2, thereby establishing itself as a new generation of small molecule fusion inhibitor against CoV infections.

## Results

### Identification of a highly conserved “E1182-L1186-L1193” motif in the HR2 domain of the SARS-CoV-2 S protein

To identify drug molecules that can destabilize the 6-HB structure, we attempted to find unique, highly conserved residues in the HR1 or HR2 domains of the SARS-CoV-2 S protein (Figure 1A) that are indispensable for 6-HB formation. For this, we analyzed the S protein sequences of all seven known human CoVs (HCoVs) including all reported variants of SARS-CoV-2, SARS-CoV, MERS-CoV, OC43, NL63, HKU1, and 229E. Sequence alignment between SARS-CoV-2 and SARS-CoV revealed high conservation in amino acid sequences for both HR1 and HR2 domains, which reduced significantly when compared with other β-HCoVs, MERS-CoV, OC43, and HKU1 and even more drastically when compared with α-HCoVs, NL63 and 229E (Figures S1A & B). Between all coronaviruses, the HR1 domain is more conserved with 7 completely conserved residues compared to 3 completely conserved residues of the HR2 domain (Figure 1B). The three conserved residues in the HR2 correspond to E1182, L1186, and L1193 of SARS-CoV-2 S. The L1186 residue remains conserved across all CoVs except hCoV-HKU1, where it is replaced with a structurally homologous isoleucine residue (Figure 1B). In addition to all HCoVs, the identity of these three residues is retained across a wide variety of non-human CoVs (Figure 1C), suggesting that these residues remain conserved throughout the course of evolution of CoVs in a wide variety of host species and hence may serve an important role in the function of the S protein.

Structural analysis of the SARS-CoV-2 S 6-HB fusion-core revealed that the E1182, L1186 and L1193 residues in HR2 participate in either intrachain or interchain hydrophobic, hydrogen, and ionic interactions with the cognate groove formed by two adjacent HR1 helices (Table S1 and Figure 1D, E, F). The negatively charged carboxylate side chain of E1182 engages in electrostatic and/or hydrogen bonding interactions with suitably placed positively charged or donor moieties within the HR1 domain of the neighboring protomer (Figure 1F, upper panel, and Table S1), while the neutral hydrocarbon side chains of L1186 and L1193 participates in multiple hydrophobic interactions with the HR1 domain of the same or neighboring spike protomers (Figure 1F, middle and lower panel, Table S1). The distances between the α-carbon atoms of E1182 and L1186 was found to be 6.3 Å, while that between L1186 and L1193 is 10.7 Å and the angle between the α-carbon atoms of E1182-L1186-L1193 is 150.2° (Figure 1G). Interestingly, the spatial organization of these three residues is completely conserved within the 6-HB structure of various other HCoVs (Figure 1G, H), which suggests that the “Ex_3_Lx_6_L” residues together constitute a highly conserved structural motif (abbreviated as “E-L-L” henceforth) responsible for the stabilization of the 6-HB.

**Table 1.** Top 5 drug candidates and their interaction profiles with the HR1-HR2 bis-peptide in their lowest binding energy conformation and in the seed conformation of the highest populated cluster. Interactions has been reported as visualized using Ligplot.

| MOLECULE/<br>CLINICAL USE | LOWEST BINDING ENERGY CONFORMATION |  |  |  | SEED CONFORMATION OF THE MOST POPULATED CLUSTER |  |  |  |
| --- | --- | --- | --- | --- | --- | --- | --- | --- |
|  | Interacting Amino acid residues |  | Binding Energy (kcal/mol) | # Conformations in the corresponding cluster | Interacting Amino acid residues |  | Binding Energy (kcal/mol) | # Conformations in the corresponding cluster |
|  | H-bonding/ ionic interactions | hydrophobic interactions |  |  | H-bonding/ ionic interactions | hydrophobic interactions |  |  |
| **Posaconazole**<br>**FDA approved**<br>Antifungal medication | **E1182**,<br>I1183,<br>S1196 | Q1180,<br>Q949,<br>L945,<br>**L1186**,<br>T941,<br>L938,<br>I934,<br>V1189,<br>**L1193**, | -6.51 | 58 | - | **E 1182**,<br>I1183,<br>Q1180, Q949,<br>L 945, **L1186**,<br>L938, I934,<br>L948, I931,<br>V1189, **L1193** | -5.92 | 84 |
| **Pirenzepine**<br>**under trial**<br>M<sub>1</sub> selective antagonist, is used in the treatment of peptic ulcers | **E1182** | L945,<br>Q1180,<br>I1183,<br>Q949 | -6.16 | 193 | **E 1182** | L 945,<br>Q1180,<br>I1183, Q949 | -6.16 | 193 |
| **Azimilide**<br>**under trial**<br>Class III antiarrhythmic drug | **E1182** | Q949,<br>Q1180,<br>I1183,<br>L945,<br>**L1186**,<br>R1185 | -6.04 | 59 | **E 1182** | Q 949,<br>Q1180,<br>I1183, L945,<br>**L1186**,<br>R1185 | -6.04 | 59 |
| **Homophenazine**<br>**Medicine possibly marketed outside USA**<br>Psychotropic drug | Q1180,<br>K1181,<br>**E1182** | Q949,<br>L945,<br>I1183,<br>**L1186** | -5.94 | 7 | S 1196 | E1195,<br>N1192,<br>F927,<br>I 934, V1189,<br>**L1193** | -4.64 | 110 |
| **BMPP**<br>**US Patent Publication (Pub. No.: US 2002/0042443 A1)**<br>Precursor to synthesize estrogen agonist/antagonist metabolites | **E1182** | N1192,<br>V1189,<br>R1185,<br>**L1186**,<br>**L1193**,<br>K1181, | -5.93 | 5 | S 1196 | **E1182**,<br>R1185,<br>**L1186**,<br>V1189,<br>**L1193**,<br>F927, L938,<br>I934, I931 | -5.25 | 83 |

### The “E-L-L” motif is critical for HR1-HR2 interaction and stability of the 6-HB structure

Molecular dynamic (MD) simulation experiments were performed to investigate the importance of the “E-L-L” motif in stabilizing the interaction between one pair of HR1-HR2 fusion-cores. Hundred nanoseconds (ns) simulation experiments were conducted using a pair of HR1-HR2 helices (residues 927-949 and 1180-1198 respectively; PDB ID: 6LXT) with either, wild type (WT) or mutant HR2 peptides harboring single, double, or triple mutations as follows: E1182A, E1182K, L1193G, E1182K-L1186G, and E1182K-L1186G-L1193G. HR1-HR2 bis-peptide with single mutations in the HR2 domain showed structural stabilities, compactness, and mobility comparable to that of the WT as evidenced by the similar root mean square deviations (RMSD) (Figure 2, A-C), radii of gyration (Rg) (Figure S2A-C), and root mean square fluctuation (RMSF) (Figure 2F, G) of the backbone atoms. In contrast, the double mutant E1182K-L1186G showed moderately high while, the triple mutant E1182K-L1186G-L1193G exhibited significantly higher RMSD (Figure 2D and E), Rg (Figure S2D and E), and RMSF values (Figure 2F and G), suggesting that the WT “E-L-L” motif is crucial for maintaining the structural integrity of the HR1-HR2 bis-peptide. It is worth noting that the small increase in the RMSD value of the E1182K (Figure 2A) mutant may indicate compromised backbone stability, due to the loss of the electrostatic/ H-bonding interactions between the HR1 and HR2 peptides.

**Figure 2.**
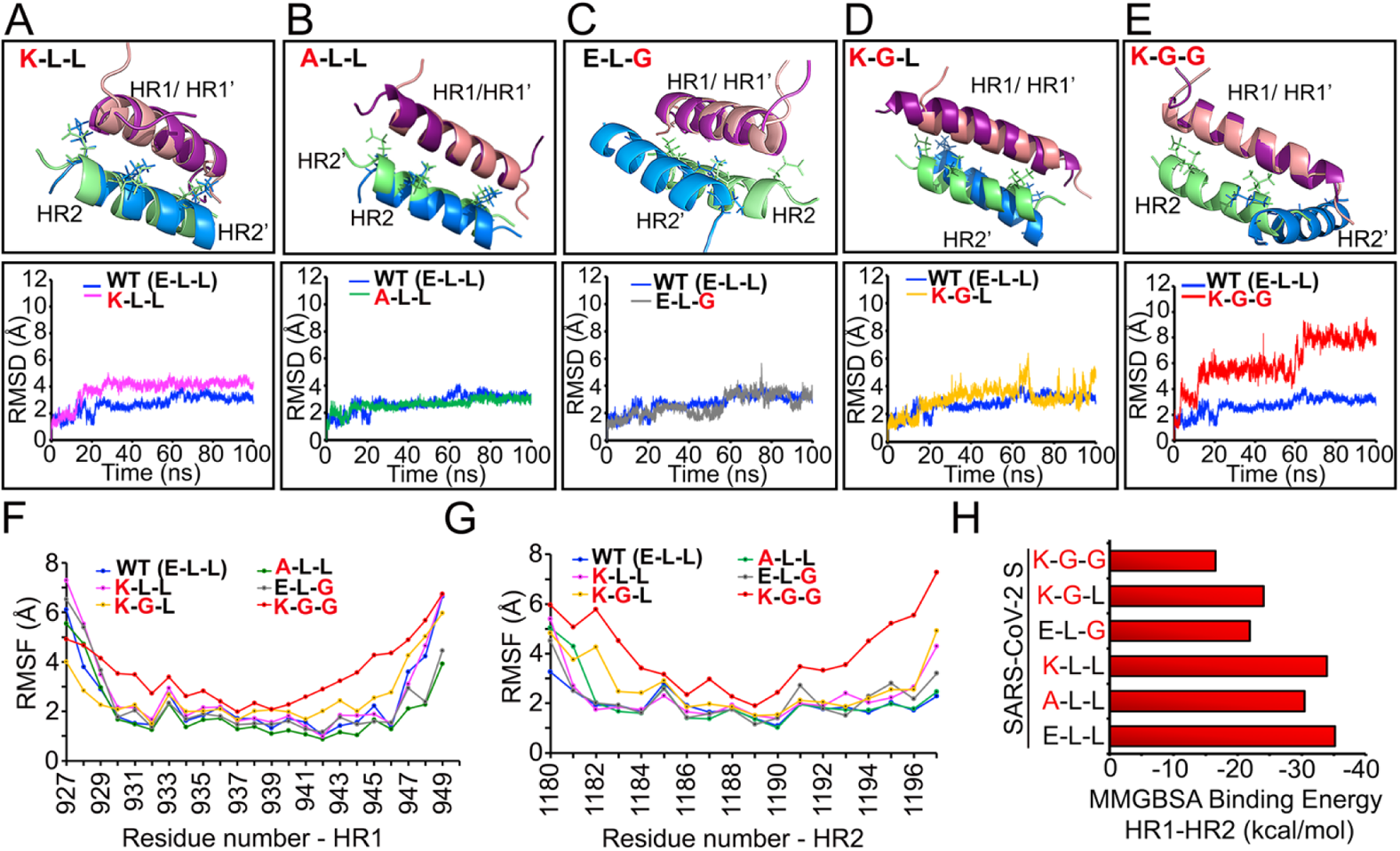
The “E-L-L” motif is critical for 6-HB stability: (A-E) MD simulation experiments showing RMSD of the backbone atoms of HR1 and HR2 helices with the WT and mutant “E-L-L” motif. Upper panels represent one of the structural snapshots of the WT (HR1/HR2) and mutant (HR1’/HR2’) bispeptides during the simulation experiment. (F & G) RMSF analyses of the atoms present within the HR1 (F) and HR2 (G) helices in the context of WT or mutant bispeptides. (H) Comparative assessment of MMGBSA binding energy for the WT and mutant dipeptides.

Estimation of the binding free energy of the WT and the different mutants by MM/GBSA method provided a more direct quantitative measurement of the interaction between these HR1 and HR2 peptides (Figure 2H). The WT bis-peptide is the most stable with ΔG value of −35.02 ± 3.83 kcal/mol, whereas E1182A and E1182K have ΔG values of −30.44 ± 2.32 kcal/mol and −33.96 ± 3.58 kcal/mol, respectively, suggesting that substitution of the glutamic acid at 1182 position exerted only marginal changes in the binding affinity between the HR1-HR2. Interestingly, the double mutant, E1182K-L1186G showed ΔG value of −24.06 ± 5.1, which is higher than that of the E1182K/ A mutants, hence indicating that the L1186 residue is also important for stabilization of the HR1-HR2 peptide complex structure. In addition, L1193 also plays a critical role in HR1-HR2 interaction as the L1193G mutant displayed ΔG value (−21.89 ± 3.43 kcal/mol) notably higher compared to other single mutants or the E1182K-L1186G double mutant. Finally, the triple mutant, E1182K-L1186G-L1193G, showed the highest free energy (−16.80 ± 3.01 kcal/mol) amongst all of the HR1-HR2 peptide complexes, reinstating the requirement of the complete “E-L-L” motif in stabilizing the HR1-HR2 interaction.

Next, we investigated the importance of the “E-L-L” motif in stabilizing the complete 6-HB structure constituted of the three pairs of the HR1-HR2 helices in the trimeric spike (PDB ID: 6LXT). MD simulation experiments were conducted for 100 ns with three pairs of HR1-HR2 helices harboring either WT (E-L-L) or mutant (K-G-G) motif in the HR2 helices. As observed from the RMSD (Figure S2F, G), the radius of gyration (Figure S2H) and RMSF (Figure S2I-N) calculation of the backbone atoms, complete alteration of the “E-L-L” motif resulted in significant perturbation of the 6-HB structure. Together, these observations strongly indicate that the “E-L-L” motif is critical for stabilizing the HR1-HR2 interaction and hence for the formation of the 6-HB/ fusion-core structure in the post-fusion form of the S protein.

### Mutations within the “E-L-L” motif impairs fusion mediated entry of SARS-CoV-2 S pseudotyped lentiviruses in ACE2 expressing cells

Next, we evaluated the importance of the identified “E-L-L” motif in S-mediated membrane fusion and viral entry using a SARS-CoV-2 S pseudovirus-based reporter assay as described previously (Crawford et al., 2020). Reporter lentiviruses pseudotyped either with SARS-CoV-2 S or the Vesicular Stomatitis Virus (VSV) G protein were used to infect either HEK293T or HEK293T-ACE2 cells and the efficiency of infection was quantified by measuring the reporter activity at 18 h of post-infection (hpi). SARS-CoV-2 S pseudotyped reporter virus showed preferential entry into the HEK293T-ACE2 cells, while the VSV G pseudotyped virus showed no such preference (Figure 3A). Subsequently, SARS-CoV-2 S pseudotyped viruses harboring single (E1182K and L1193G), double (E1182K-L1186G), or triple (L1182K-L1186G-L1193G) mutations within the “E-L-L” motif were compared with WT S pseudotyped viruses for their ability to infect HEK293T-ACE2 cells (Figure 3B). Mutations either at E1182 or at L1193 caused 40% and 60% reduction in reporter activity respectively, confirming the importance of these two residues in the optimum activity of the S protein. In addition, L1186 was also found to be crucial as pseudoviruses harboring E1182K-L1186G mutations showed a 65% decrease in reporter activity. Finally, replacing all three residues of the motif (E1182K-L1186G-L1193G) resulted in an 85% reduction in reporter activity (Figure 3B), confirming that complete integrity of the “E-L-L” motif is crucial for S-mediated entry of pseudoviruses in target cells. All the mutants were expressed and were proteolytically cleaved at the S1-S2 junction with efficiencies comparable to the WT S protein (Figure 3C).

**Figure 3.**
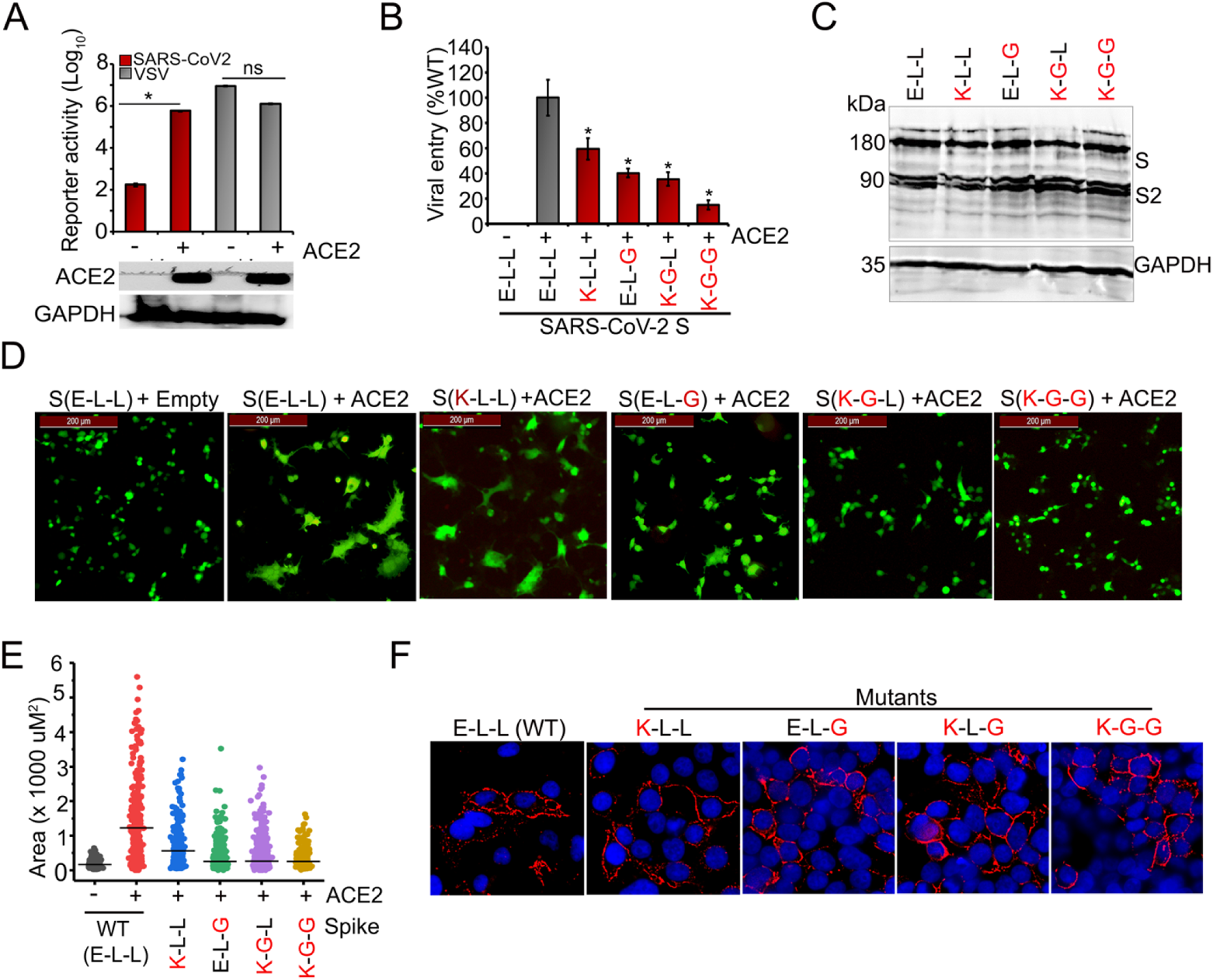
Mutation in the “E-L-L” motif blocks fusion mediated viral entry. (A) SARS-CoV-2 S or VSV G protein pseudotyped lentiviruses viruses were used to infect either HEK293T or HEK293T-ACE2 cells. Luciferase activities were monitored as a measure of virus entry. (B) SARS-CoV-2 S protein (WT or mutants) pseudotyped viruses were tested for their ability to infect HEK293T-ACE2 cells. (C) WB analysis of the WT or mutant S proteins, expressed in HEK293T cells, using S specific antibody (D) HEK293T overexpressing SARS-CoV-2 S (WT or mutants) and eGFP were co-cultured with HEK293T-ACE2 cells for 12h and syncytium formation was monitored (E) Area of the GFP positive cells was measured from ten independent fields to determine the mean syncytium area using Image J. (F) HEK293T cells were transfected with WT or mutant S protein-expressing plasmids and at 48 h post-transfection surface expression of the S proteins were evaluated using an anti-spike antibody.

After synthesis, a fraction of the S protein reaches the cell surface and is embedded into the cell membrane where they trigger multinucleated syncytia formation between infected and uninfected cells (Buchrieser et al., 2020). Hence, we investigated the importance of the “E-L-L” motif in spike-mediated membrane fusion and syncytium formation. HEK293T cells overexpressing SARS-CoV-2 S along with eGFP (HEK293T-S+GFP) were co-cultured with HEK293T-ACE2 cells for 24h before fixing and imaging. Cells expressing WT S protein generated multinucleated GFP positive syncytia, while, cells expressing the E1182K mutant showed moderate reduction and the E1193G and E1182K-L1186G mutants exhibited a severe reduction (Figure 3D, E, and Figure S3) in syncytium formation. More significantly, cells expressing the triple mutant, E1182-L1186G-L1193G were completely defective in membrane fusion as syncytia formation was rarely observed in these cells (Figure 3D, E, and Figure S3). We also confirmed that all the S proteins were expressed on the cell surface at similar levels through surface immunostaining (Figure 3F), indicating that these mutations in the HR2 domain do not impede S protein processing and maturation. These data along with the MD simulation observations unambiguously establish that the “E-L-L” motif plays an indispensable role in S-mediated membrane fusion by stabilizing the HR1-HR2 interaction and 6-HB structure formation.

### *In silico* screening for drugs targeting the highly conserved “E-L-L” motif

The high conservation of the “E-L-L” motif and its importance in fusion-mediated viral entry drove us to explore it as a potential drug target for developing a broad-spectrum fusion inhibitor against coronaviruses. We mined standard chemical databases (PubChem, DrugBank Online, and Zinc15) and selected 25 potential candidates (22 FDA approved or under trial drugs and 3 previously reported small molecules, Figure 4, S4), that have stereo-electronic complementarity with the conserved “E-L-L” motif and under physiological conditions, possess either both or at least any one of the following structural features – (i) bear positively charged or polar side chains capable of interacting with E1182 either through salt-bridge or hydrogen bonding interactions, and (ii) structural components like aryl rings and/or hydrocarbon (acyclic/cyclic) side chains capable of interacting with L1186 and L1193 through hydrophobic interactions.

**Figure 4.**
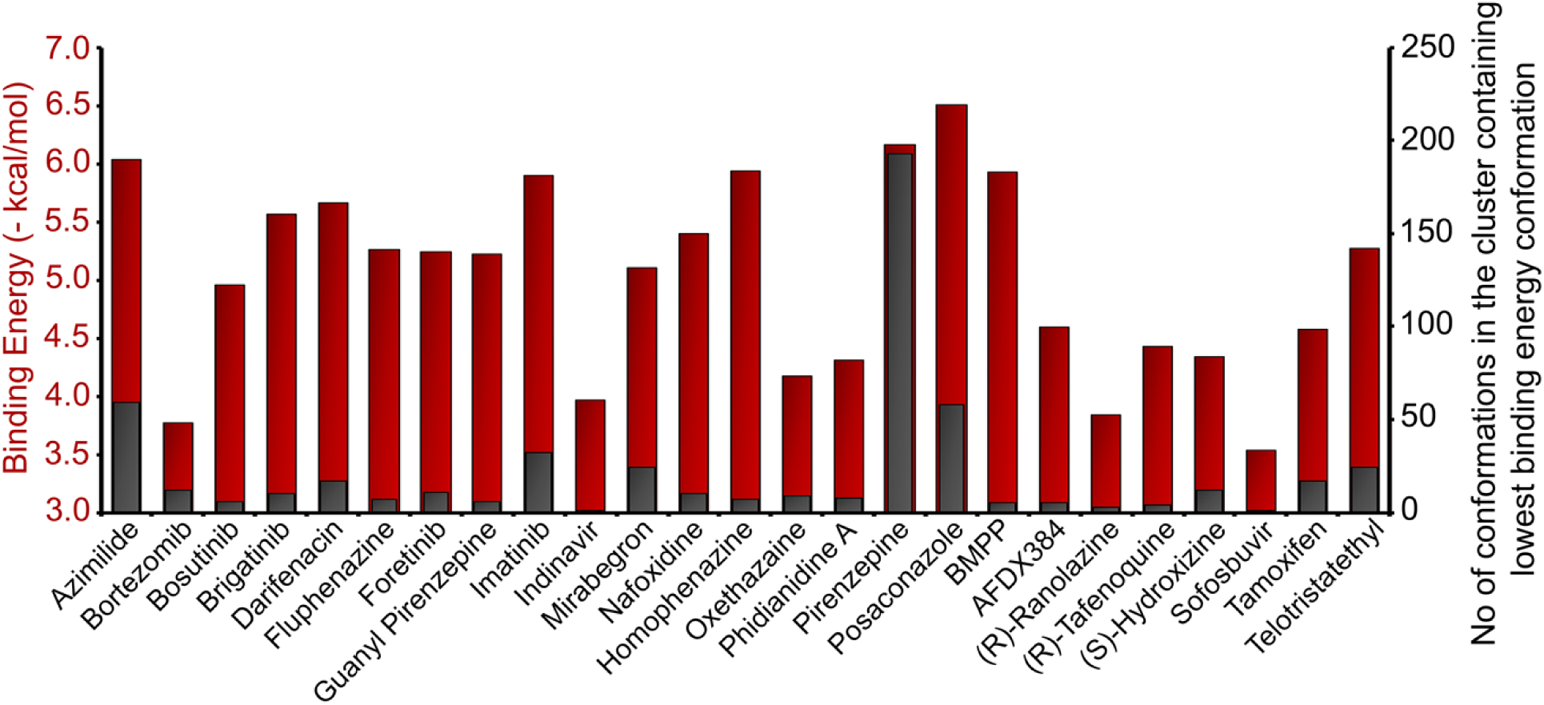
In silico screening of drug molecules targeting “E-L-L” motif. Comparative plot of the lowest binding energies and number of conformations present in the cluster containing the lowest binding energy conformation corresponding to each of the docked molecules.

Molecular docking studies (Balaji et al., 2013; Novikov & Chilov, 2009) were carried out with a single pair of HR1-HR2 fusion-core helices as a receptor with the AutoDock grid (implemented through AutoDock 4.2) (Morris et al., 2009) set around the conserved “E-L-L” motif (Figure 5, S6, S7). We hypothesized that binding of the drug with at least any one of the HR1-HR2 pairs would sterically hinder its association with other HR1-HR2 helices and therefore block the fusion process by destabilizing the 6-HB structure. To get more statistically accurate results, 500 docked conformations were generated for each of the 25 docked ligands, which were then grouped into different clusters based upon their conformational similarity (Varela-Salinas et al., 2017). Each cluster contains one lowest energy seed conformation along with other structurally similar conformations (RMSD tolerance of 2.0Å), all of which binds to a single pocket within the receptor. Thus, the number of docked conformations belonging to a specific cluster could be directly correlated with the probability of the ligand binding to a particular pocket. Using this analysis, we classified individual drugs based on three criteria; lowest binding energy; number of conformations in the cluster containing lowest binding energy conformation, and finally, ability to interact with the conserved “E-L-L” motif (Table 1 and S2). Figure S5 represents a three-dimensional plot for comparative analysis of the 25 drug molecules based upon all three criteria while Figure 4 represents a two-dimensional plot based upon the first two criteria.

**Figure 5.**
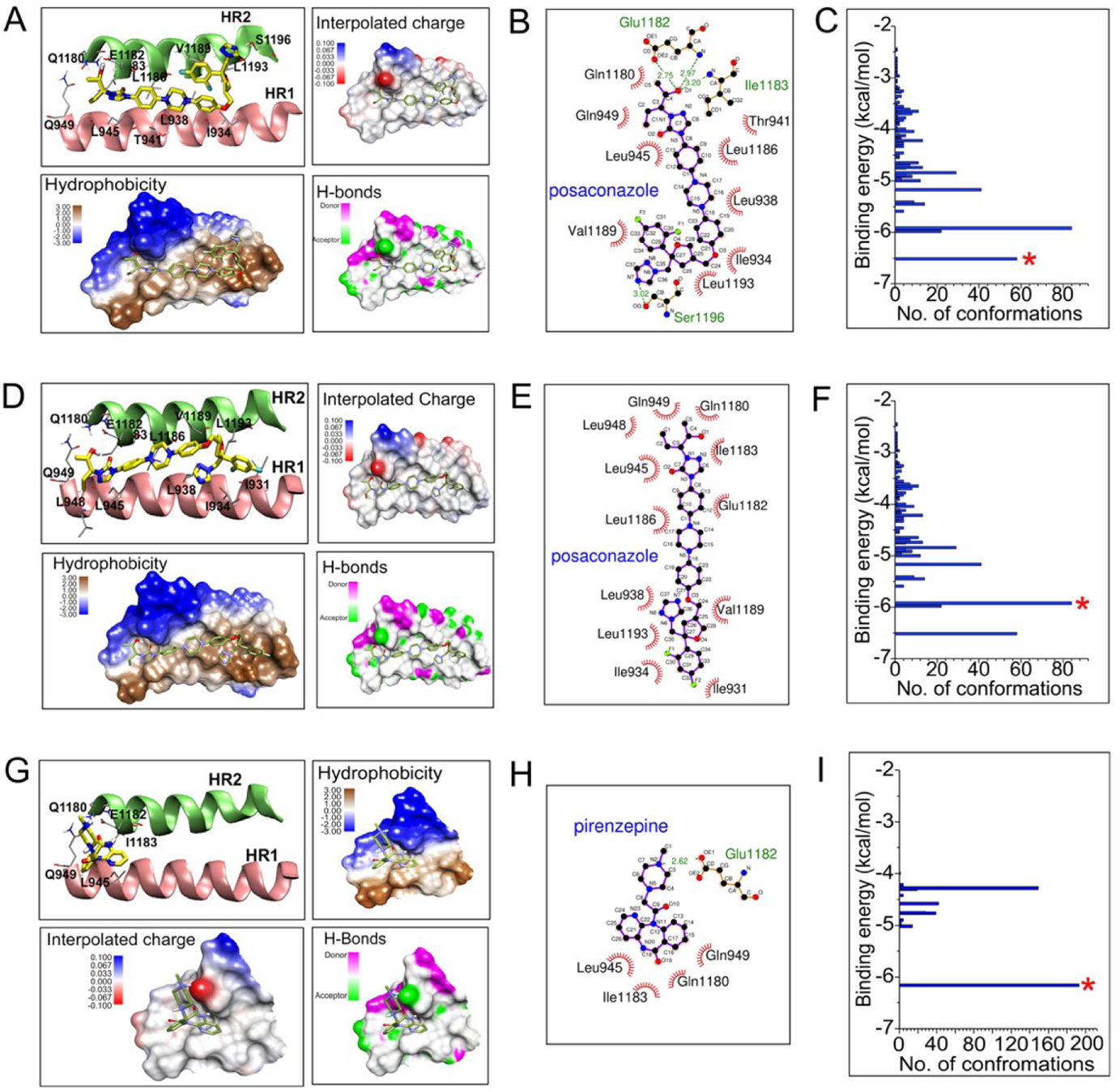
Interaction profiles of Posaconazole and pirenzepine to the HR1-HR2 helices. Images A-C corresponds to the lowest binding energy conformation of posaconazole; D-F corresponds to the seed conformation of the most populated cluster for posaconazole; G-I corresponds to the lowest binding energy conformation of pirenzepine. A, D, and G represent the binding pocket for each of the corresponding ligand conformation (HR1 and HR2 helix shown as cartoon image, ligand molecule, and the amino acid side chains shown in stick model); hydrophobicity, hydrogen bond donor/acceptor and interpolated charges on the protein surface for each corresponding ligand conformation have also been mapped. B, E, and H are the ligplot images showing the corresponding ligand-protein interaction in two-dimension. C, F and I, represents the cluster analysis with the asterisk marks denoting the particular cluster to which the corresponding ligand conformation belongs.

### Posaconazole shows high affinity towards the “E-L-L” motif-containing binding pocket

To identify the drug with the highest potential of inhibiting the HR1-HR2 interaction, we ranked the drugs based upon their binding energy and focused on the top five candidates – posaconazole, pirenzepine, azimilide, homophenazine, and BMPP (Figure 4, Table 1), and analysed their interaction profile with the “E-L-L” motif. Posaconazole (Figure 5 A-C) and BMPP (Figure S6 I-L), in their lowest binding energy conformation, interacted with the largest number of residues spanning across the length of the HR1-HR2 helices, including all three residues of the “E-L-L” motif. In contrast, azimilide and homophenazine interacted with fewer residues, which included E1182 and L1186 (Figure S6 A-D, and E-H) while, pirenzepine interacted selectively with E1182 along with a few more surrounding residues (Figure 5 G-I). It is important to note that, posaconazole and BMPP showed similar interaction profiles both in their highest populated cluster and in the cluster containing lowest binding energy conformation (Figure 5 A-C & D-F, Table 1), while for pirenzepine and azimilide, the cluster containing the lowest binding energy conformation remained highest populated (Figure 5 G-I and Figure S6 A-C, Table 1). For homophenazime, the seed conformation of the highest populated cluster showed significantly lesser binding affinity than the lowest energy one and interacted only with L1193 of the “E-L-L” motif (Figure S6 I-K and Table 1). Based on these analyses, we focused our attention on posaconazole and pirenzepine; the first one showed the lowest binding energy and interacts with the entire E-L-L motif while the second one, although interacting preferentially with only E1182, showed considerably high binding affinity (only next to posaconazole’s lowest energy conformation) and possesses the highest number of conformations in the lowest binding energy cluster.

Posaconazole, in its most stable docked conformation (*cluster population of 58*) as well as in the seed conformation for the highest populated cluster (*cluster population of 84*) (Figure 5 C and F), predominantly occupied the hydrophobic groove formed by the HR1-HR2 domains (Figure 5 A and D). In its most stable docked conformation, the −OH (O1) group of posaconazole participates in two hydrogen-bonding interactions with E1182 and one hydrogen bonding with I1183 (Figure 5 A, B table S3). The triazole nitrogen (N7) forms hydrogen bonding with –OH from S1196 (table S3). The other structural components like the aryl rings and hydrocarbon linkers were found to contribute towards hydrophobic stabilization of the ligand-protein complex (Figure 5 B). In the seed conformation of the most populated cluster (Figure 5 D, E), posaconazole interacted exclusively through hydrophobic interactions and almost completely occupied the hydrophobic groove created by the HR1, HR2 domain. Pirenzepine, on the other hand, in its most stable docked conformation (*with a cluster population of 193*) (Figure 5 G-I) showed strong binding affinity only towards the HR2 N-terminal and HR1 C-terminal regions. Perhaps lack of extended hydrophobic core and predominance of polar functional groups disfavored its entry into the hydrophobic groove (Figure 5 G). The quaternary positively charged N (N2) showed strong salt-bridge/ hydrogen bonding interaction with the negatively charged oxygen from the carboxylate residue in E1182 (Figure 5 H).

### Posaconazole inhibits SARS-CoV-2 S pseudotyped lentivirus entry, specifically targeting the “E-L-L” motif

Having determined posaconazole and pirenzepine to be the two most suitable candidates for binding to the “E-L-L” motif, and destabilizing the HR1-HR2 interaction, we tested the ability of these two drugs in blocking SARS-CoV-2 S pseudotyped lentivirus entry into ACE2 expressing cells. SARS-CoV-2 S pseudotyped lentiviruses were pretreated with increasing concentrations of either posaconazole or pirenzepine before infecting HEK293T-ACE2 cells, which were then further incubated in the respective drug concentration until harvest. Treatment with posaconazole resulted in a dose-dependent decrease in SARS-CoV-2 pseudovirus entry with more than 90% reduction in reporter gene expression at a concentration of 10µM (Figure 6A), whereas pirenzepine did not have any significant effect (Figure 6B). The half effective inhibitory concentration (IC_50_) of posaconazole was estimated to be 3.37µM (Figure 6C). However, posaconazole did not inhibit the entry of control VSV G pseudotyped viruses (Figure 6A), which enters cells via endocytosis followed by endosomal fusion, suggesting that the drug specifically blocks SARS-CoV-2 entry, possibly by selectively targeting the “E-L-L” motif of the S protein. We also showed that posaconazole does not inhibit the entry and infection caused by other enveloped viruses like influenza A/H1N1 virus (Figure 6D) or herpes simplex virus 1 (HSV1) (Figure 6E), hence establishing its specificity in blocking the entry of SARS-CoV-2. Interestingly, pirenzepine, although showed similar binding affinity (6.16 kcal/mol) to posaconazole (6.51 kcal/mol) *in silico* (Table 1), failed to impact entry of SARS-CoV-2 S (Figure 6B), possibly because it specifically interacts only with the E1182 and not with the whole “E-L-L” motif. Both the drugs were non-toxic at the concentrations used for the pseudovirus reporter assay (Figure S8).

**Figure 6.**
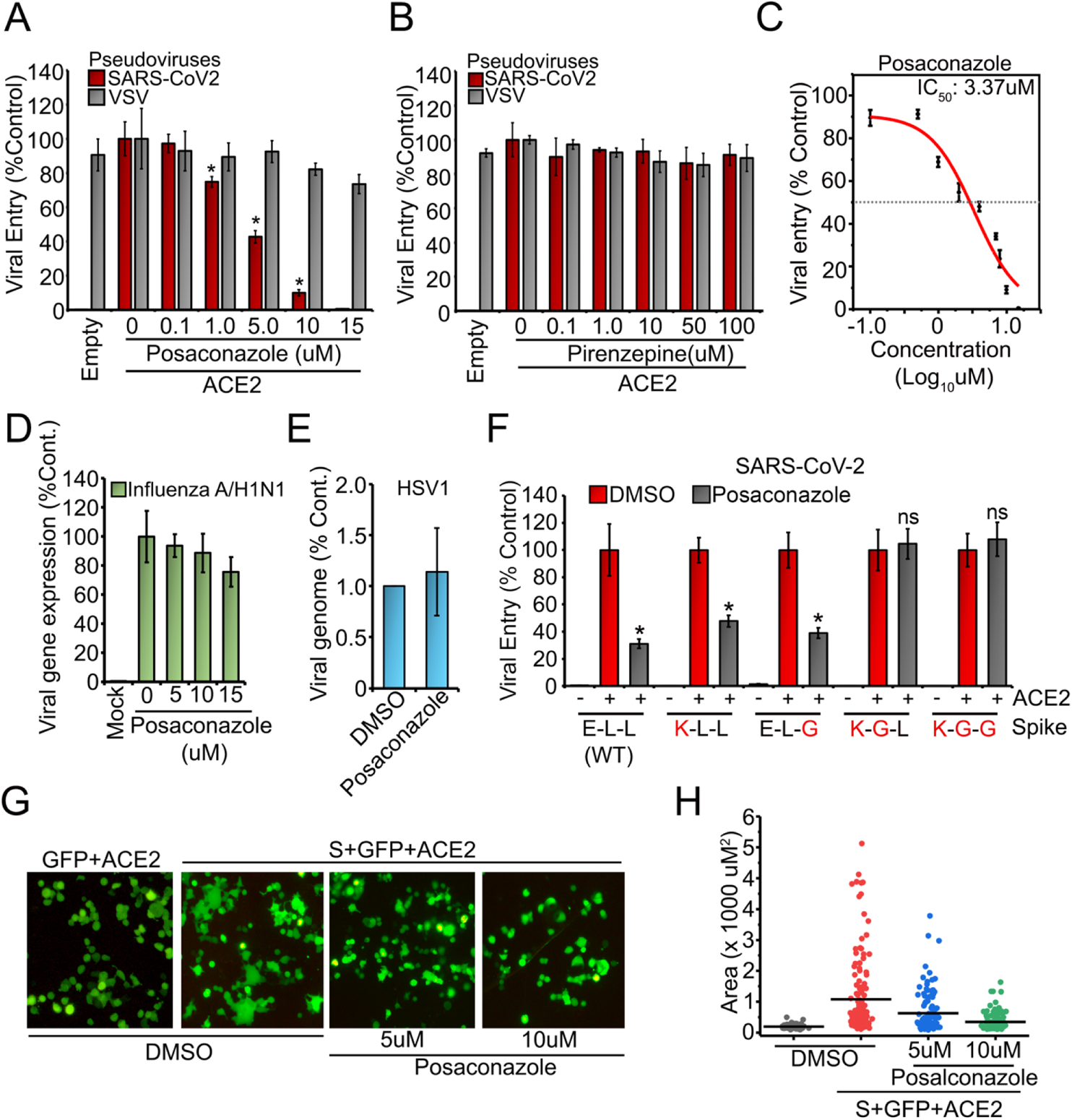
Posaconazole inhibits SARS-CoV-2 S pseudotyped lentivirus entry and syncytium formation. (A, B) SARS-CoV-2 S or VSV G pseudotyped reporter viruses and were pretreated with DMSO or with different concentrations of the indicated drugs and used to infect either HEK293T (Empty) or HEK293T-ACE2 cells in presence of the drug. Reporter activity was measured at 18 h and plotted as a relative percentage to the DMSO treated sets. (C) IC_50_ of posaconazole against the SARS-CoV-2 S pseudotyped viruses was calculated by fitting reporter assay data in the four-parameter nonlinear equation in origin lab software. (D) MDCK cells were infected with the influenza A (WSN/H1N1/1933) nano-Luc reporter virus pretreated with DMSO or with different concentrations of posaconazole and reporter activity was measured at 8 hpi. (E) Vero cells were infected with HSV1 pretreated with DMSO or with 8µM posaconazole and viral genome was quantified at 2hpiusing qPCR. (F) WT or mutant SARS-CoV-2 S pseudoviruses were treated with posaconazole before, during, and after infecting HEK293T-ACE2 cells with them. Reporter activity was measured and plotted as a relative percentage of the DMSO treatment for each set (WT and mutant viruses). (G) Cells expressing SARS-CoV-2 S and GFP were treated with different concentrations of posaconazole or with DMSO before and during co-culturing them with HEK293T-ACE2 cells for syncytium formation assay. (H) The area of the GFP positive cells was measured from five independent fields and the mean area plotted using Image J software. *=p < 0.001 (unpaired student’s t-test). Images shown are representative of three independent repeats.

To unambiguously establish the selectivity of posaconazole towards the “E-L-L” motif, we tested its efficacy against viruses pseudotyped with the SARS-CoV-2 S protein harboring single, double, or triple mutations within the “E-L-L” motif. As described earlier (Figure 3), these mutant pseudoviruses although shows variable degrees of attenuation in their infectivity owing to the defect associated with the membrane fusion potential of the mutant S proteins, can still mediate entry in HEK293T-ACE2 cells. Pseudoviruses harboring single S mutations, E1182K and L1193G, showed almost similar sensitivity towards posaconazole compared to the WT (Figure 6F). In contrast, pseudoviruses harboring double or triple mutation within the “E-L-L” motif became highly resistant towards posaconazole. Clearly, substituting two of the three residues of the “E-L-L” motif makes the SARS-CoV-2 S protein completely resistant towards the drug, hence establishing the high selectivity of posaconazole towards this motif.

Given that the “E-L-L” motif is indispensable for membrane fusion and posaconazole shows high selectivity towards this motif, we next tested the ability of posaconazole to inhibit membrane fusion by syncytia formation assay. Pretreatment of HEK293T-S+GFP cells with increasing concentrations of posaconazole and their subsequent co-culturing with HEK293T-ACE2 cells resulted in significant reduction in syncytium formation in comparison to the control set (Figure 6G, H and S9), confirming that posaconazole can specifically block membrane fusion triggered by SARS-CoV-2 S protein. Together, the data presented so far suggest that posaconazole specifically inhibits SASR-CoV-2 S mediated entry, by blocking the fusion of viral and host membranes, specifically targeting the highly conserved “E-L-L” motif present in the HR2 domain of the S protein.

### Posaconazole efficiently inhibits S mediated entry of newer SARS-CoV-2 variants of concern (VOC)

Of late, the delta (B.1.617.2) and kappa (B.1.617.1) variants of SARS-CoV-2 have been a matter of immense concern (https://www.who.int/en/activities/tracking-SARS-CoV-2-variants/, 2022; Pascarella et al., 2021). The delta variant has been declared as one of the main VOC associated with higher infectivity, hospitalization, reinfection, and infection in vaccinated individuals (Mlcochova et al., 2021). More recently, the omicron (B.1.1.529) VOC was identified with a staggering 30 mutations in its S gene and has been predicted to have even higher infectivity than the delta variant (Chen et al., 2022). Interestingly, the “E-L-L” motif and all of the residues across HR2 and HR1 helices that participate in interaction with posaconazole are highly conserved across all different variants including the delta, kappa, and omicron VOCs (Figure 7A, S10), suggesting that posaconazole should be effective against these more virulent variants of SARS-CoV-2. As evidenced from the data, posaconazole exhibited strong inhibitory effects on the entry of the pseudoviruses harboring S protein from D614G variant, the dominant clade of SARS-CoV-2 (Plante et al., 2021), the kappa (B.1.617.1), delta (B.1.617.2) and the omicron (B.1.1.529) variants which is comparable to that of the original Wuhan strain (Genbank NC_045512) (Figure 7B), hence confirming its efficacy against newer and deadlier variants of SARS-CoV-2.

**Figure 7.**
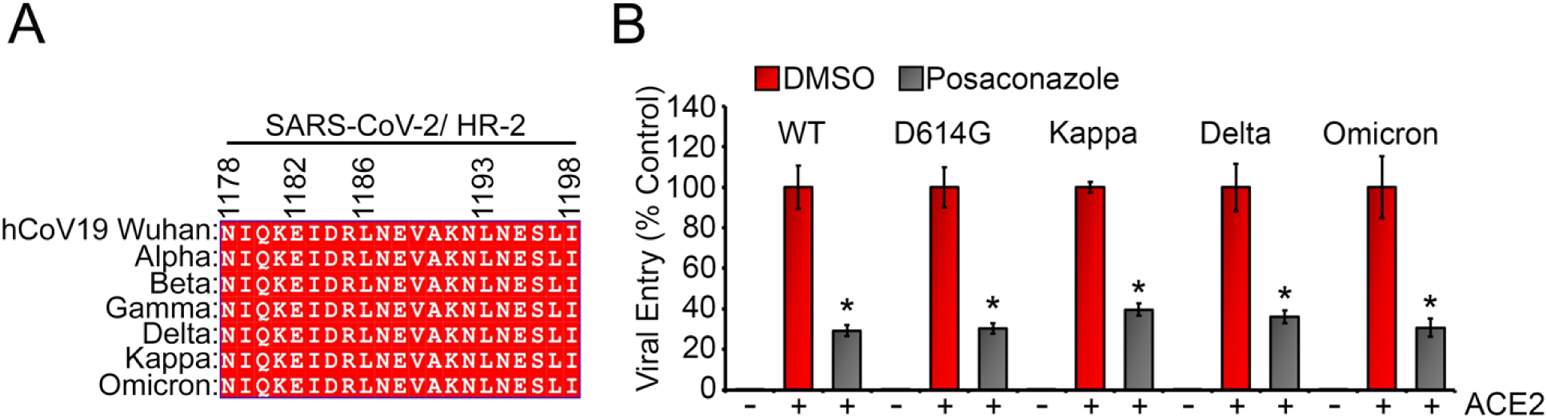
Posaconazole blocks the entry of different variants of SARS-CoV-2 pseudoviruses. (A) Amino acid alignment of the posaconazole interacting region of HR2 from major variants of concerns (VoC) as obtained from the GISAID database. (B) HEK293T-ACE2 cells were infected with pseudoviruses expressing S proteins of either WT SARS-CoV-2 (Wuhan strain) or major VoC’s (D614G, Delta, Kappa, Omicron), treated with DMSO or with posaconazole before, during, and after infection. Reporter activity was plotted as a relative percentage to the DMSO control for each set. *=p < 0.001 (unpaired student’s t-test). Images shown are representative of three independent repeats.

### Posaconazole inhibits SARS-CoV-2 infection through blocking S-mediated membrane fusion and subsequent release of viral genomic RNA in Caco-2 cells

Next, we tested the efficacy of posaconazole as a fusion inhibitor on infectious SARS-CoV-2 virus. Using a fluorescently labelled recombinant virus based on the 2019-nCoV/USA-WA1/2020 strain, SARS-CoV-2-mNG (Xie et al., 2020), we evaluated three different concentrations of the drug, 4, 8, and 12 µM, for anti-SARS-CoV-2 activity. Caco-2 cells, pretreated with the respective concentrations of posaconazole were infected with SARS-CoV-2-mNG, also pretreated with the corresponding concentrations of the drug. 48 hpi, we observed a dose-dependent decrease in mNG signal (Figure 8A, B, and Figure S11) indicating efficient inhibition of SARS-CoV-2 infection. We followed up this experiment with a plaque assay under similar conditions but using non-recombinant WT SARS-CoV-2 and observed greater than 1 log reduction of virus titer at all 3 concentrations tested, with 8 and 12 µM showing the greatest but similar inhibition of the virus (Figure 8C and Figure S12A). Posaconazole was non-toxic at these concentrations (Figure S8D), and we selected 8 µM for all downstream experiments. Posaconazole also inhibited the infection of more recent variants of SARS-CoV-2, including the Beta variant (also known as South African variant) and the Delta variant at efficiencies equal to the WT variant (Figure 8D and Figure S12B). Furthermore, we evaluated the fusion inhibition potential of posaconazole on SARS-CoV-2-mNG infected Caco-2 cells by visualizing and measuring the syncytia formed after 48 hpi (Figure 8E, F and Figure S13). As evident, posaconazole treatment drastically reduced syncytium size confirming that it inhibits membrane fusion between adjacent cells.

**Figure 8.**
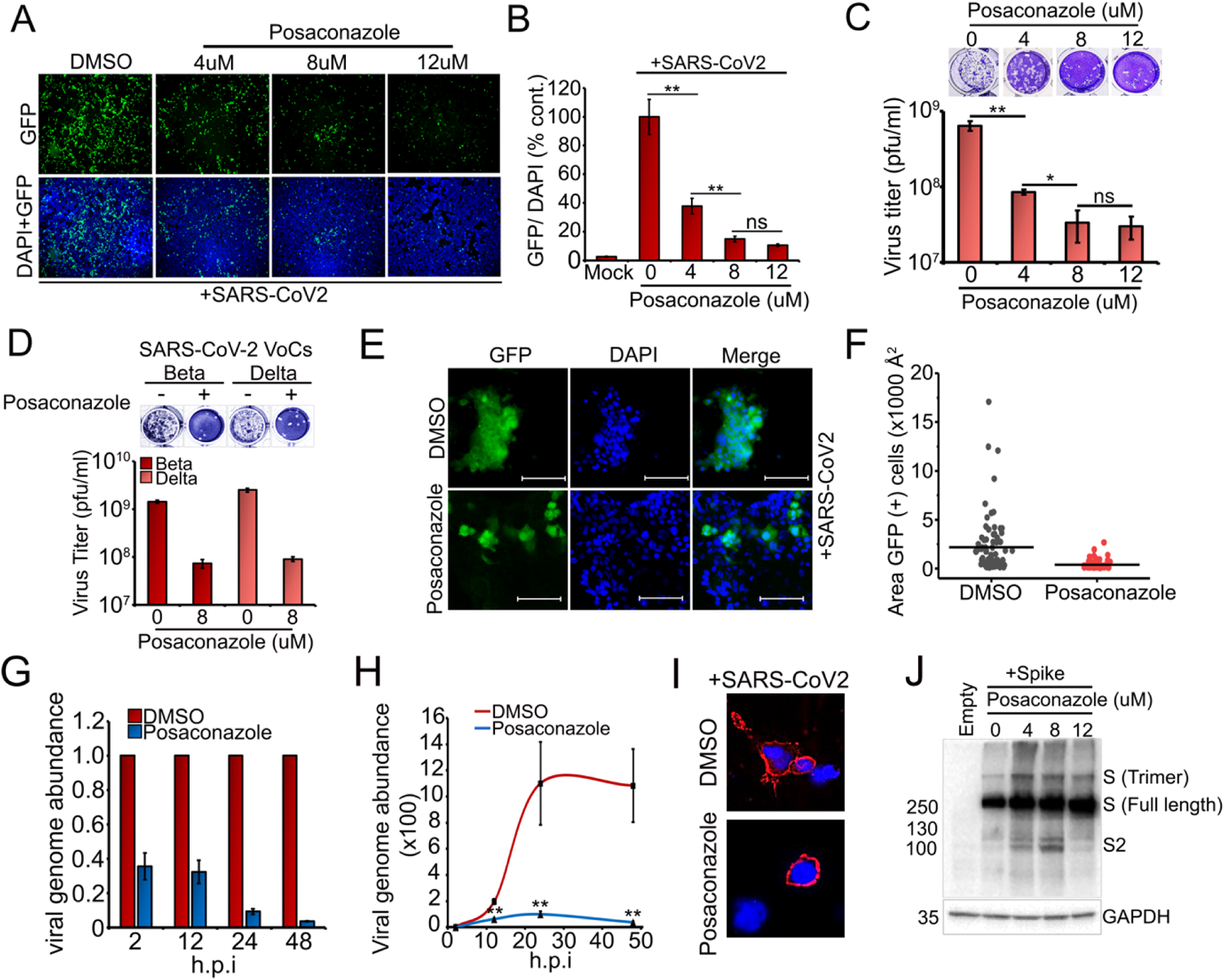
Inhibitory effects of posaconazole on infectious SARS-CoV-2 virus. (A) Caco-2 cells pretreated for 1h with DMSO or the indicated concentrations of posaconazole were infected with SARS-CoV-2-mNG virus (multiplicity of infection, MOI of 0.1) also pretreated with corresponding concentrations of the drug. 48 hpi, mNG fluoresence were visualized and quantitated using ImageJ (B). mNG signals were normalized against DAPI signals, and plotted relative to DMSO treated samples. (C, D) Caco-2 cells were infected either with WT SARS-CoV-2 (C) or with the beta or delta variants (D) pretreated with DMSO or posaconazole (8µM) and after 48hpi, the infected supernatants were titrated using plaque assay. (E) Caco-2 cells, pretreated with posaconazole (8µM) were infected with pretreated SARS-CoV-2-mNG (MOI 0.1) and visualized at 40X for syncytia formation. (F) The area of each syncytium was measured using ImageJ and plotted. (G) Caco-2 cells pretreated with DMSO or posaconazole (8 µM) and were infected with pretreated SARS-CoV-2. Total RNA extracted at indicated times, were evaluated by qRT-PCR. The data are expressed as fold changes of the viral N RNA levels normalized to RNaseP control relative to the corresponding DMSO treated samples. (H) Similar experiment as in G, but expressed as relative to the initial inoculation amount (at 2 hpi). (I) Surface immunofluorescence assayfor the S protein was performed on Caco-2 cells infected with WT SARS-CoV-2 in presence of DMSO or posaconazole (8 µM). (J) WB of the S protein 72h after infection of Caco-2 cells with WT SARS-CoV-2 in presence of DMSO or different concentrations of posaconazole. All data shown are averages of the results of at least three independent experiments SD. Images shown are representative of all independent repeats.

As successful fusion of viral and host cell membranes leads to the release of the viral genome into the host cell, we evaluated the effect of posaconazole upon the accumulation of SARS-CoV-2 genomic RNA at different time points post-infection. Cells were infected with SARS-CoV-2 with or without posaconazole treatment and qRT-PCR was performed to quantify the amount of viral genomic RNA at different time points post-infection. When normalized to its corresponding mock treatment, posaconazole resulted in a gradual decrease in viral RNA abundance with time (Figure 8G) with more than 95% reduction at 48 hpi. Interestingly, at 2-12 hpi, we observed a 64% reduction in the abundance of viral RNA, indicating that posaconazole inhibited the initial infection of the virus (2 hpi) as well as subsequent rounds of progeny virus release and infection of neighboring cells (24 and 48 hpi). When normalized to the initial inoculum of viral RNA (2 hpi), the DMSO treated cells exhibited an exponential increase in viral RNA up to about 30 hpi, where after, it plateaued (Figure 8H). Posaconazole treated cells under similar conditions exhibited greatly reduced virus replication kinetics confirming its robust anti-viral potential (Figure 8H). We also confirmed that posaconazole neither affected the migration of S protein to the cell membrane (Figure 8I and Figure S14), nor did it affect S protein synthesis and/or stability (Figure 8J).

Finally, we performed time of treatment assays to study the efficacy and mechanism of action of posaconazole. As illustrated in figure 9A, for a prophylactic setting, viruses and the cells were (a) pre-incubated with posaconazole for 1h, which were then used for infection in absence of the drug, or (b) treated during the infection followed by washing off the drug. We also included sets where cells were treated with posaconazole (c) only post-infection, or (d) both during and after infection. In another set (e), both virus and cells were treated with posaconazole at ‘all times’, pre, during and post infection. As observed (Figure 9B, C and Figure S15), post-treatment of the drug alone had a much-diminished effect on reducing infection compared to the other time of addition experiments. Moreover, treatment only during infection had an almost similar outcome as the ‘all-time’ treated sets. Both pre+during and during+post treatments had comparable inhibitions to the ‘all-time’ treated samples. This confirms that posaconazole functions as a fusion inhibitor and its presence during infection is enough to exhibit a robust anti-viral effect. Additionally, post-infection treatment of the cells with posaconazole also showed a significant effect, possibly rooted through its effect on blocking subsequent rounds of infection of neighboring cells. Together, these data confirm that posaconazole act as a fusion inhibitor that can inhibit SARS-CoV-2 at the entry stage of infection and also suggests that it has both, prophylactic, as well as, therapeutic potentials.

**Figure 9.**
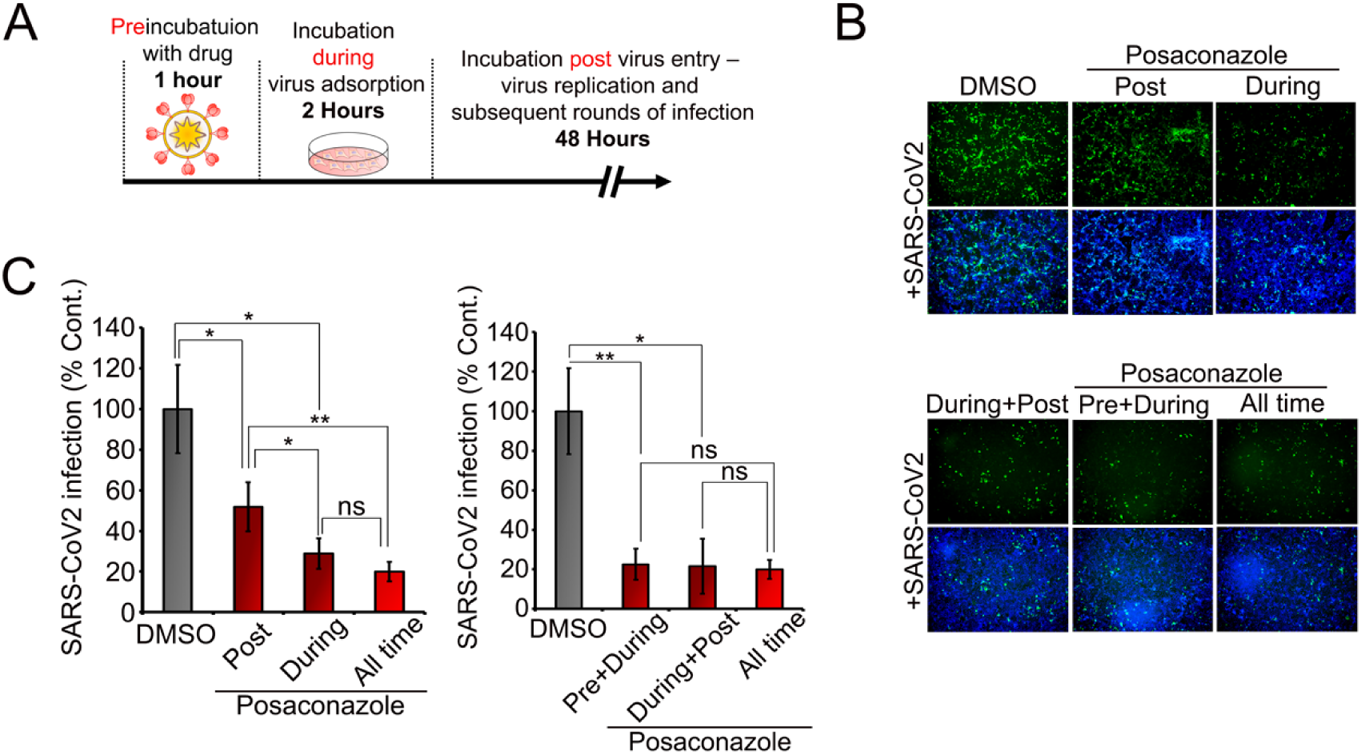
Posaconazole blocks SARS-CoV-2 infection at early stage. (A) Schematic representation of the time of addition assay for posaconazole (8 µM) on Caco-2 cells infected with SARS-CoV-2-mNG (MOI 0.1). (B, C) mNG was visualized and quantitated by counting the mNG positive cells and normalized to the total DAPI signal in that field. All data shown are averages of the results of at least three independent experiments ± SD. *=p < 0.05; **=p < 0.01; ***=p < 0.001, ****= p <0.0001 (unpaired student’s t-test). Images shown are representative of all independent repeats.

## Discussion

The rapid evolution of the SARS-CoV-2 resulting in recurrent surges of infection by different VOCs and its concomitant effect on global health and economy (https://www.who.int/en/activities/tracking-SARS-CoV-2-variants/, 2022) calls for the development of novel therapeutic strategies which will be able to combat both, the current and future variants. In this report, we describe the identification of a novel drug target in the viral spike glycoprotein and its subsequent targeting using a safe, pharmacologically approved molecule to develop an effective entry inhibitor against SARS-CoV-2.

Fusion inhibitors are of great potential to block the entry of enveloped viruses, which require fusion of the viral and host cell membranes to release their genetic material into the cytoplasm. Diverse families of viruses, including *Retroviridae*, *Orthomyxoviridae*, *Paramyxoviridae*, *Filoviridae,* and *Coronaviridae*, possessing Class I fusion proteins, share common structural mechanism for triggering membrane fusion. Interaction between three HR1 and HR2 domains of the trimeric fusion protein, leads to the formation of the 6-HB, which is critical for bringing the viral and host membranes in close proximity, hence driving their fusion (Bosch et al., 2003; Chambers et al., 1990; Y. Huang et al., 2020; Weissenhorn et al., 1999; White et al., 2008). This makes the 6-HB structure an attractive drug target for blocking membrane fusion and viral entry (Bird et al., 2014; Dongen et al., 2019; Kandeel et al., 2021; Lan et al., 2021; Mzoughi et al., 2019; Outlaw et al., 2020; S Xia et al., 2019; Shuai Xia et al., 2018; Yang et al., 2020; Y Zhu et al., 2020; Yuanmei Zhu et al., 2021). However, till date, the 6-HB structure has been mostly targeted using long peptides (35-50 amino acids) mimicking either HR1 or HR2 helices (Bosch et al., 2004; Kesteleyn et al., 2017; S Xia et al., 2019; X. Zhu et al., 2015; Yuanmei Zhu et al., 2021). While these peptides show high efficacy in inhibiting virus replication, their widespread implementation as antiviral drugs will be challenging owing to the inherent limitations of peptide-based therapeutics (Fosgerau & Hoffmann, 2015; Otvos & Wade, 2014). On the other hand, bioavailable small molecule fusion inhibitors, which are both easier to produce and deliver than peptides, have been disappointingly scarce and only a handful have been described for IAV and HIV (Dongen et al., 2019; Zhoua et al., 2011) and more recently against CoVs (Hu et al., 2021; Park et al., 2022; Yang et al., 2020).

Successful targeting of viral infection with small molecules requires identification of a suitable “molecular target” possessing the following characteristics, (i) it should represent a structural element or a sequential motif or a combination of both, which would offer a suitable interaction interface for the small molecule; (ii) the target region should be critical for the activity of the viral protein and (iii) it should be minimally tolerant towards evolutionary changes acquired through antigenic drifts or shifts. Considering these parameters, in this study, we identified a highly conserved motif, E1182-L1186-L1193 present within the HR2 helix of SARS-CoV-2 S protein, whose side chains participate in essential molecular interactions indispensable for the stability of the 6-HB (Figure 1 B, C, E and F). MD simulation analysis suggested that mutation of these residues perturb the HR1-HR2 interaction and thus, alter the overall stability of the 6-HB fusion-core structure (Figure 2 A-H). This was supported by our mutational analysis, which showed that alteration of either single or multiple amino acids within the “E-L-L” motif makes the S protein attenuated towards fusion and entry of SARS-CoV-2 pseudoviruses in ACE2 overexpressing cells (Figure 3 B, D and E). Together, these data confirmed importance of the “E-L-L” motif in structure and activity of S protein and thus establishes itself as a novel molecular target for the discovery of a new generation of fusion inhibitors against CoVs.

We undertook a multidimensional *in silico* screening strategy to identify potential FDA-approved or under trial drugs capable of targeting this “E-L-L” motif. For this, we classified the drugs based on their binding energy towards the “E-L-L” motif, the number of conformations that are closely related to the lowest binding energy conformation (RMSD=2Å) and their potential to interact with individual amino acids constituting the motif. Posaconazole, emerged as the best possible candidate with significantly high binding affinities in two of the highest populated clusters and numerous molecular interactions spanning across the entire “E-L-L” motif. While posaconazole shows strong fusion inhibitory activity against the WT SARS-CoV-2 S protein (Figure 6A) more than one mutation within the “E-L-L” motif makes the protein completely resistance towards the same (Figure 6F), hence, substantiating the high specificity of the drug towards its target. Though the appearance of such mutations might seem to be of concern for possible appearance of posaconazole-resistant variants, the high conservation of the “E-L-L” motif across all reported variants of SARS-CoV-2 and amongst other CoVs point towards the high genetic/evolutionary barriers that these viruses have to overcome in order to develop such resistance. Additionally, high degree of amino acid conservations suggest that the SARS-CoVs(SARS-CoV-2 and SARS-CoV) are least tolerant towards alterations in their HR2 domain (Shuai Xia, Zhu, et al., 2020), hence establishing it as an ideal target for broad-spectrum fusion inhibitors.

Posaconazole is an azole antifungal drug approved by the US FDA in 2006 (Schlossberg & Samuel, 2017). In our studies, it exhibited high efficacy in inhibiting entry and replication of SARS-CoV-2 and its VoCs. Syncytia formation in infected lungs cells has been recognized as a hallmark of severe stages of SARS-CoV-2 infection with strong implications in the pathogenic outcome of the patient (Braga et al., 2021). Our data confirmed that posaconazole specifically inhibits S-mediated membrane fusion, syncytium formation and subsequent release of viral genome in host cells, substantiating its anti-fusion role. We show that posaconazole blocks the virus at early stages of infection, making it superior to the currently available drugs that target RdRp activity and hence block the later stages of infection (Buonaguro et al., 2020). Therefore, an exciting proposition towards a novel anti-SARS-CoV-2 therapy can be the use of posaconazole as a prophylactic nasal spray to inhibit early colonization of the virus in the upper respiratory tract and subsequent administration of oral suspension of the drug as a therapeutic to prevent or treat lower respiratory tract infection. The pharmacokinetics of posaconazole is well studied where a recommended 400 mg twice daily dosage results in mean peak drug plasma concentrations (Cmax) of 4.15 µg ml−1, or 6 µM (Lipp, 2010), which is comparable to the effective cell-culture concentration of the drug we observed in our studies. Posaconazole showed a favorable safety profile, with only ⩽2% of patients exhibiting side effects when administered in a dose of 400 mg/day for 24 weeks (Catanzaro et al., 2007). Most interestingly, posaconazole is under clinical trial for treating COVID19 associated pulmonary aspergillosis (Atul Patel et al., 2021; Hatzl et al., 2020), hence establishing it as a safe option to treat both SARS-CoV-2 and associated secondary fungal infections. Recent *in-silico* analysis also point towards posaconazole’s role as a SARS-CoV-2 helicase inhibitor (Abidi et al., 2021), which is however yet to be experimentally validated.

Together, our study provides the mechanism and rationale for repurposing posaconazole as a potential anti-SARS-CoV-2 drug. Easy and ready availability, low toxicity, and well-known pharmacokinetics of posaconazole along with its proven efficacy against different variants of SARS-CoV-2 will support initiation of fast-track clinical trials for rapid dissemination of the drug as the next generation antiviral therapy to treat COVID19 disease.

## Materials and methods

### Plasmids

Plasmid HDM-IDTSpike-fixK, expressing a codon-optimized spike protein from SARS-CoV-2, Wuhan-Hu-1 strain (Genbank NC_045512) under a CMV promoter (BEI catalog number NR-52514), was used in this study. All site-directed mutagenesis was performed on this codon-optimized spike clone as the WT form using the QuikChange II Site-directed mutagenesis kit (Agilent), following manufacturer’s instructions. Primers used are provided in Supplementary Table 4 and all mutations were verified using Sanger’s sequencing. Plasmid pHAGE2-EF1aInt-ACE2-WT, expressing the human ACE2 gene (BEI #NR-52514) was used for all ACE2 overexpression studies. Plasmids pCMV VSV-G, expressing the VSV-G (addgene#138479) (Gee et al., n.d.), pcDNA3.3-SARS2-B.1.617.2 SARS-CoV-2, expressing the spike protein of the Delta variant (addgene#172320), pcDNA3.3-SARS2-B.1.617.1, expressing the spike protein of the Kappa variant (addgene#172319)(Cho et al., 2021), and SARS-CoV-2 Omicron Strain S gene Human codon_pcDNA3.1(+) expressing the spike protein of the Omicron variant (GenScript# MC_0101274) were used as indicated.

### Cell lines

Human embryonic kidney (HEK) 293T cells (ATCC #CRL-3216), Madin Darby Canine Kidney (MDCK) cells (ATCC #CCL-34), and African green monkey kidney epithelial (Vero) cells (ATCC #CCL-81) were maintained in Dulbecco’s modified Eagle’s medium (DMEM) supplemented with 10% fetal bovine serum (FBS) along with penicillin and streptomycin antibiotics (Gibco) at 37°C and 5% CO_2_. Human colon epithelial cells (Caco-2) cells (ATCC #HTB-37), were maintained in Minimum Essential Medium (MEM) media supplemented with 20% fetal bovine serum, Penicillin-Streptomycin, 1x non-essential amino acid solution, and 10 mM sodium pyruvate at 37°C and 5% CO_2_.

### Antibodies and Drugs

Goat polyclonal antibody against human ACE2 (AF933) and rabbit polyclonal antibody against SARS CoV-2 Spike protein (NB100-56578) were obtained from Novus Biologicals (USA). Posaconazole (SML2287) and Pirenzepine dihydrochloride (P7412) were purchased from Sigma Aldrich (USA).

### Viruses

SARS-CoV-2 Isolates USA-WA1/2020, South Africa/KRISP-K005325/2020 and USA/MD-HP05647/2021 (lineage B.1.617.2; Delta variant) were obtained from BEI Resources (BEI #NR52281, #NR-54009, and #NR-55672, respectively). SARS-CoV-2-mNG, a recombinant SARS-CoV-2 virus having a genomic backbone of the 2019-nCoV/USA_WA1/2020 strain and harboring the mNeonGreen (mNG) gene in the ORF7 was a kind gift from Dr. PEI-Yong Shi from the University of Texas Medical Branch, Galveston, TX, USA (Xie et al., 2020). All SARS-CoV-2 stocks were produced and purified from supernatants of Vero-E6 ACE2 cells, infected with different variants of SARS-CoV-2 or SARS-CoV-2-mNG virus, at MOI of 0.1 in 10 ml MEM supplemented with 5% FBS. 72 hpi, virus-containing supernatants were harvested, centrifuged (500g, 10 min), filtered (0.45 μm), mixed with 10% SPG buffer (ATCC #MD9692), aliquoted and stored at −80°C. For quantification of virus titers, viral stocks were serially diluted (10-fold) in serum-free medium and inoculated on 1×10^5^ Vero E6-ACE2 cells, followed by determination of plaques forming units (PFU) after 72 h of infection on confluent Vero-E6-ACE2 cells. All replication-competent SARS-CoV-2 experiments were performed in a biosafety level 3 laboratory (BSL-3) at the University of South Florida. The influenza A/WSN/33 (H1N1), PA-2A-Swap-Nluc (PASTN), reporter influenza virus expressing nano-luciferase, is a kind gift from Prof. Andrew Mehle, University of Wisconsin Madison (Tran et al., 2013) and was propagated and purified from MDCK cells. Herpes Simplex Virus 1 (HSV-1 KOS) was propagated and purified from Vero cells.

### Bioinformatics and structural analysis

Multiple sequence alignment was performed in Bioedit software using clustalW multiple alignment function. Amino acid sequences of different human infecting Corona virus spike proteins were obtained from NCBI virus database (accession no: QXI74535.1, AYV99761.1, QCQ29075.1, AGT51481.1, ADN03339.1, QRK03775.1, AGT51338.1). HR1 and HR2 regions of the specific proteins were aligned using the clustalW multiple sequence alignment function in Bioedit. Representative image was generated using ESPript 3.0 website (https://espript.ibcp.fr/ESPript/cgi-bin/ESPript.cgi). Multiple sequence alignment data was used to calculate amino acid identity matrix in Bioedit for HR1 and HR2 regions. Sequences of HR1 and HR2 region of different variants of SARS-CoV2 were also aligned using clustalW multiple sequence function in Bioedit software. Conservation of E-L-L motif was checked by searching “Ex_3_Lx_6_L” sequence in different animal coronavirus HR2 region in Bioedit. Identified regions are aligned and logoplots were generated using WebLogo website (https://weblogo.berkeley.edu/logo.cgi). Structural alignment was performed in Pymol using specific PDB files of post-fusion spike protein structures of different human infecting coronaviruses (PDB id: 6LXT,1WYY,4NJL,5YL9 and 2IEQ). Intrachain and interchain protein-protein interactions were calculated using PIC webserver (http://pic.mbu.iisc.ernet.in) with the default distance parameters.

### Molecular Dynamics Simulations

The HR1-HR2 peptide complexes harboring either WT or different mutant variants of the HR2, were subjected to molecular dynamics simulations for a period of 100 ns using Gromacs software (Abraham et al., 2015). For MD simulation of the six Helix-bundle structure, three chains, A, B, and C from the crystal structure of the SARS-CoV-2 S protein (PDB ID: 6LXT) were used. Together they contained a complete 6-HB structure covered by the interactions between all three sets of HR1-HR2 interactions. Mutations in the helix-bundle chains were incorporated using the Chimera program. CHARMM36 force filed (J. Huang & MacKerell Jr, 2013) and TIP3P water model (Jorgensen et al., 1983) were used for the MD simulations in the explicit solvent conditions. In order to mimic physiological concentration at a neutralized condition, 0.15 M salt concentration was used. Number of Sodium and Chloride ions to be added before the simulation was calculated using packmol software (Martínez et al., 2009). A rectangular box was created where the boundary of the box was at least 10 Å away from the solute molecules. Steepest descent algorithm was used to energy minimize the solvated protein structures with maximum force limit of 1000 kJ mol^-1^ nm^-1^. For equilibration, 100 ps simulations were carried out using NVT and NPT ensembles at temperature 300K and pressure 1 bar, respectively. Gromacs tools were utilized to analyze the trajectories from 100 ns simulations. MMGBSA analyses were carried out using gmx_MMPBSA software (Valdés-Tresanco et al., 2021), which is an extended work of MMGBSA.py for Gromacs trajectories (Miller et al., 2012). For MMGBSA analyses, 200 snapshots were used from the last 10 ns of the MD simulations at an interval of 50 ps. MMGBSA values were then plotted using Origin software.

### Screening of small molecules; Molecular Docking

Autodock4.2 was used to predict the binding modes and binding energies for the drug candidates. Ligand PDB structures were built up using Chem 3D Pro or procured as SDF file data bases like PubChem, Drugbank online etc. Protonation states for the ligands at physiological pH were considered using UCSF Chimera (Pettersen et al., 2004) interface. The structures were further geometry optimized using Avogadro (Hanwell et al., 2012) via MMFF94 force field. Ligand PDB files were edited in Autodock tools (ADT) by setting the root, torsions and Gasteiger charges were assigned, finally saved as PDBQT files. The fusion core HR1-HR2 helices was used as the receptor. After the separation of the coordinates of heteroatoms, water molecules were removed, polar hydrogens were added. Kollman charges were calculated with AutoDock Tools (ADT). A grid box of 48 × 36 × 90 points and a spacing of 0.375 Å, having center at X= −2.335, Y=1.035, Z=-28.732, was used with an objective to sufficiently cover the conserved motif E1182, L1186 and L1193. Grid potential maps were calculated using module AutoGrid 4.0. During docking, the number of Genetic Algorithm runs, maximum number of generations, maximum number of evaluations, size of individuals in the population are set to 500, 27000, 2500000 and 150, respectively. The ranking of the poses was done based on binding energies, ability to interact with the conserved residues, nature of interactions and finally cluster analysis. The structures were visualized using MGL Tools-1.5.6, Discovery Studio 2019 Client and VMD (Humphrey et al., 1996). The 2D-interactions were interpreted from the image obtained through LigPlot v2.2.4(Wallace et al., 1995).

### SARS-CoV-2 S and VSVG pseudotyped lentivirus generation

Lentivirus virus particles pseudotyped with full-length WT and mutant SARS-CoV-2 S protein were generated by transfecting HEK293T cells as described by Katharine et. al (Crawford et al., 2020). Briefly, the lentiviral backbone plasmid pHAGE-CMV-Luc2-IRES-ZsGreen-W (BEI #NR-52516) that uses a CMV promoter to express luciferase followed by an IRES and ZsGreen, was co-transfected with plasmids pHDM-Hgpm2, expressing the HIV-1 gag and pol (BEI #NR-52517), pRC-CMV-rev1b, expressing the HIV-1 rev (BEI #NR-52519), pHDM-tat1b, expressing the HIV-1 Tat (BEI #NR NR-52518), and HDM-IDTSpike-fixK or its mutants at a 4.6:1:1:1:1.6 ratio. The media was changed 18 h after transfection, followed by collection of the lentivirus containing supernatant, 60 hours later. The supernatant was clarified by passing through a 0.45 um filter and used freshly. For lentivirus virus particles pseudotyped with VSV-G protein, a similar protocol was followed, replacing HDM-IDTSpike-fixK with pCMV VSV-G.

### Pseudovirus based entry assay

For pseudovirus-based entry assay, HEK293T cells were transiently transfected with lentivirus expression plasmid of human ACE2 receptors. At 24 hours of post-transfection, hACE2 overexpressing cells were infected with reporter lentiviruses pseudotyped with SARS-CoV-2 S (WT or mutants) or VSV-G proteins as their only surface glycoprotein. Luciferase gene expression was measured at 24 hpi by using a Glomax 20/20 luminometer (Promega, USA) using the luciferase assay reagent (E1501, Promega, USA). HEK293T cells transfected with the respective empty vectors were used as negative controls. For mutant spikes, luciferase activities were plotted as relative percentage of the WT SARS-CoV-2 pseudoviruses. Three independent experiments, each having three replicates were performed.

### Analyzing the effect of drugs on pseudovirus entry

Reporter lentiviruses pseudotyped with either SARS-CoV-2 spike protein (S) or Vesicular Stomatitis Virus (VSV) glycoprotein (G) at its surface or otherwise mentioned were treated with different concentrations of either posaconazole or pirenzepine for 30 min at RT. HEK 293T cells expressing hACE2 were infected with treated viral inoculum for 1h at 37^0^C in presence of the respective concentrations of the drug. The cells were further replenished with fresh media with the same concentration of the drugs. Infected cells were harvested at 18 hpi and luciferase activity was measured using a luminometer-Glomax 20/20 (Promega, USA).

### Cell-cell fusion assay

HEK293T cells were transfected with plasmids expressing hACE2 or co-transfected with plasmids expressing SARS-COV-2 S protein (WT or mutants as mentioned) and Green Fluorescence Protein (GFP). 36 h post-transfection both the cell lines were trysinised and co-cultured in 1:1 ratio for 12 h followed by fixing permialization and subsequent imaging using fluorescence microscope (Leica Microsystems) and subsequent analysis by ImageJ software.

### MTT assay

Cellular cytotoxicity of the drugs were determined by MTT assay as previously mentioned by Mosmann et al. (Mosmann, 1983). Briefly, cells seeded in a 96-well plate at a density of 30,000 cells per well or otherwise mentioned were subjected to different concentrations of the drugs for 24 h at 37°C in 5% CO_2_. Thereafter, 100μL of MTT reagent (5mg/mL, SRL) in PBS was added to the cells and incubated for 3 h at 37 °C. Finally, the precipitated formazan crystals were dissolved in 100μL of dimethyl sulfoxide (DMSO) (Sigma, USA) and the absorbance was measured at 595 nm using an Epoch 2 microplate reader (BioTek Instruments).

### Nano-Luc Reporter Assay

Luciferase activity of nanoluciferase influenza A/H1N1/WSN/1933 reporter virus was determined as previously described by Tran et al (Tran et al., 2013). Briefly, the influenza A reporter virus was preincubated with Posaconazole for 30 min at RT and subsequently used to infect MDCK cells seeded in a 96-well plate. The reporter activity of the was measured at 8hpi using Nano-Glo luciferase assay system (Promega) using a Glomax 20/20 luminometer.

### Cell-cell fusion assay

HEK293T cells transfected with lentiviral expression plasmids expressing hACE2 or empty vector using Lipofectamine 3000 reagent (Thermo Fisher Scientific, USA). In parallel, HEK293T cells were co-transfected with plasmids expressing SARS-COV-2 S protein (WT or mutants as mentioned) and Green Fluorescence Protein (GFP). 36 hours post-transfection spike and GPF co-expressing cells were trypnised, and seeded over the hACE2 expressing cells in 1:1 ratio and incubated at 37^0^C for 12 h. The cells were fixed with 3% formaldehyde and permeabilized with 0.1% Triton X-100 and imaged in Leica Fluorescence microscope (Leica Microsystems). Images were analyzed by ImageJ software.

### SARS-CoV-2 infection and Drug treatment

All SARS-CoV-2 infections were performed using the same passage SARS-CoV-2 variants or SARS-CoV-2-mNG virus stocks. Caco-2 cells were pre-treated with the appropriate posaconazole concentration for 12h. Viral stocks were appropriately diluted to the indicated MOI in serum-free medium and also pre-incubated with the corresponding posaconazole concentration for 2h. After pre-incubation, cells were washed two times in serum-free medium and inoculated with the appropriate drug-treated virus, and 2h after infection, cells were washed with complete medium, and infection allowed to proceed for the indicated time points in DMEM supplemented with 2.5% FBS and the corresponding concentration of posaconazole. Subsequently, at appropriate times post-infection, cells or supernatants were harvested and analyzed using appropriate techniques.

### Surface immunofluorescence

For detection of the WT and mutant spike protein localizing to the surface of the cell, HEK293T cells were seeded on 8-chamber glass-bottomed slides and transfected with appropriate plasmids. 48 h post-transfection, cells were washed and treated with freshly prepared 0.1% paraformaldehyde and incubated for 10 minutes at 4°C. Cells were then washed two times with PBS and incubated with an anti-spike antibody diluted (1:200) in PBS for 1 h at 37°C. Subsequently, cells were washed three times with PBS for 5 minutes each and incubated with an appropriate secondary antibody (1:500) for 30 minutes at RT in the dark. Finally, the cells were washed 3 times with PBS and mounted using an anti-fade reagent. Imaging was performed on a Keyence BX700 microscope. For evaluation of the effect of posaconazole on the surface localization of the spike protein, Caco-2 cells were pre-treated with the drug or the vehicle control for 12 h and then infected with SARS-CoV-2 in the presence of the drug. 48 hpi, cells were fixed and processed as described above.

### RNA extraction and qRT-PCR

Total RNA was isolated from appropriately treated cells using the RNeasy mini-kit (Qiagen #74106) following manufacturer’s instructions. On-column DNase digestion was performed by using an RNase-free DNase set (Qiagen #79254). 1 μg RNA was reverse transcribed by using the High-Capacity cDNA reverse transcription kit (Applied Biosystems #4368814), according to the manufacturer’s instructions. For qPCR, the synthesized cDNA was diluted 1:20 and used as a template with Power SYBR Green PCR Master Mix (Applied Biosystems #4367659) on an ABI Prism 7500 detection system (Applied Biosystems) using the SARS-CoV-2 N gene-specific primer set, Forward: CACATTGGCACCCGCAATC and Reverse: GAGGAACGAGAAGAGGCTTG. All mRNA levels were normalized to RNaseP mRNA levels and calculated as the delta-delta threshold cycle (ΔΔCT).

### Western blot analysis of SARS-CoV-2 S protein

The expression of spike glycoprotein of SARS-CoV-2 in HEK293T cells were detected by western blot. Briefly, cells were transfected with plasmids expressing either SARS-COV-2 S protein WT or mutants (E1182K, L1193G, E1182K/L1193G, L1182K/L1186G/L1193G) by using Calcium phosphate transfection kit (Takara-Bio). 24 h post transfection, the cells were lysed using RIPA buffer (50 mM Tris-HCl [pH 7.5], 150 mM NaCl, 2mM EDTA, 1% NP-40, 0.5% deoxycholate, 0.1% SDS) supplemented and equal amount of total protein samples were separated in an 8% SDS PAGE. Proteins were transferred to PVDF membrane, probed using anti SARS CoV-2 S antibody (Novus Biologicals, USA) and visualized with Chemiluminescent Reagent (Pierce).

### Statistical analysis

All experiments were performed in triplicates and each data was repeated at least three times. Graphs are plotted using Microsoft Excel and Origin and represented as mean standard deviations (n = 3). Results were compared by performing a two-tailed Student’s t test. Significance is defined as P < 0.05, and statistical significance is indicated with an asterisk (*). *P < 0.001 were considered statistically significant.

## Acknowledgements

We sincerely thank Dr. PEI-Yong Shi from the University of Texas Medical Branch, TX, USA for providing the SARS-CoV-2-mNG. The Nano-Luciferase reporter influenza virus was a kind gift from Prof. Andrew Mehle, University of Wisconsin Madison, WI, USA. We gratefully thank BEI resources NIAID, and Addgene for SARS-CoV-2 S gene related plasmids, lentivirus plasmids and other constructs as mentioned in the materials and methods section. We acknowledge funding support from DBT, Ramalingaswami re-entry fellowship (BT/RLF/Re-entry/02/2015), SERB, Early Career Research Award (ECR/2017/001896), MHRD, “Scheme for Transformational and Advanced Research in Science” (STARS/APR2019/BS/369/FS (Project ID: 369)), ICMR (Grant No. VIR/COVID-19/19/2021/ECD-I); awarded to AM, Public Health Service grant R01 CA180758 to BC, Veterans Affairs Merit Review grant BX005490 and Research Career Scientist Awards IK6BX004212 and IK6BX003778 awarded to SM and SSM.

## Authors contribution

Conceptualization: AM, AR and DB.

Methodology & Design: AM, AR, IDJ, PB, SB, DB and GB

Investigation: IDJ, AR, PB, SB, KM, AM, GB, SD, AG, DB, SA, SS and ARM.

Supervision: AM, AR, SM

Writing—original draft: AM, AR, IDJ, PB and DB

Writing—review & editing: AM, AR, IDJ, PB, GB, SB, AKD, AB, BC, SM, SSM Fund acquisition-AM, BC, SM & SSM.

## Data and materials availability

All data are available in the main text or the supplementary materials.

## Supplementary information

**Figure S1.**
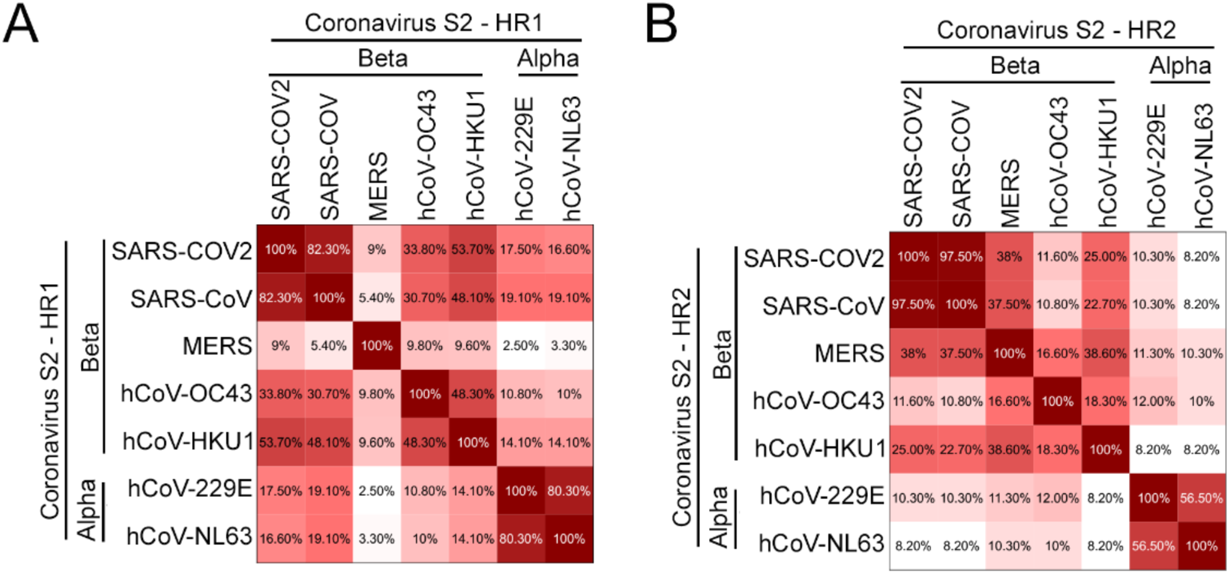
Amino acid identity matrix of (A) HR1 and (B) HR2 region of spike protein among different human infecting coronavirus.

**Figure S2.**
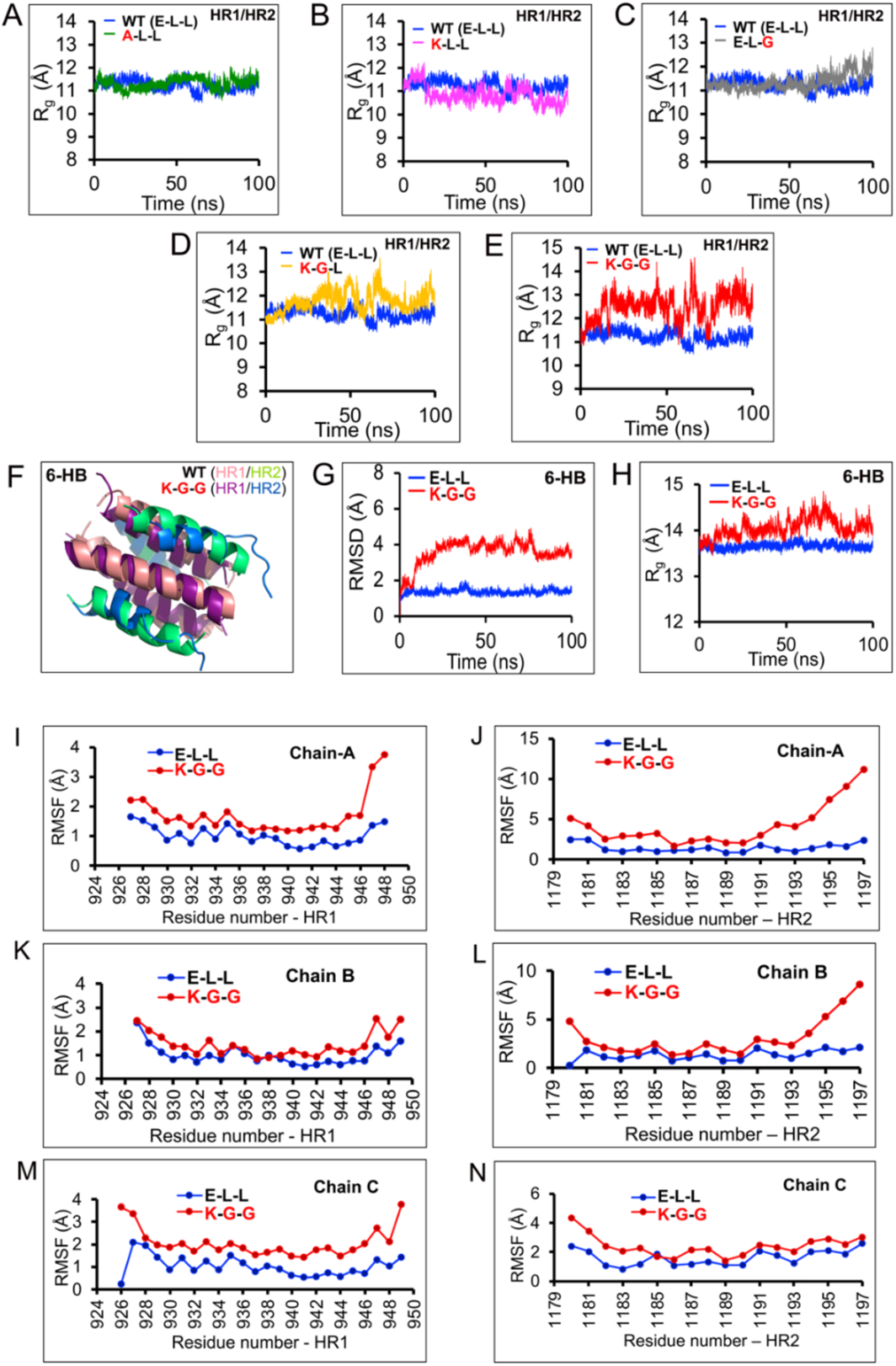
MD simulation analysis of HR1-HR2 peptide complex. (A-E) Radius of gyration (Rg) analysis of the trajectories obtained for the mutants with respect to the WT HR1-HR2 dipeptide. Dipeptide harboring triple mutant E1182K-L1186G-L1193G) is the least compact among all the model dipeptides. (F-N) MD simulation experiment of the 6-HB structure either harboring the WT “E-L-L” motif or the triple mutant “K-G-G”. (F) Structural snapshot of the WT (HR1 pale orange and HR2 green) or triple mutant (HR1 magenta and HR2 blue) 6-HB structure during the MD simulation experiment. (G) RMSD and (H) Rg analysis of the WT or mutant 6-HB structures showing significant perturbation in case of the mutant. (I-N) RMSF analyses of the atoms present within the HR1 (I, K, M) and HR2 (J, L, N) helices when considered in the context of WT or mutant dipeptides situated within the 6-HB structure.

**Figure S3.**
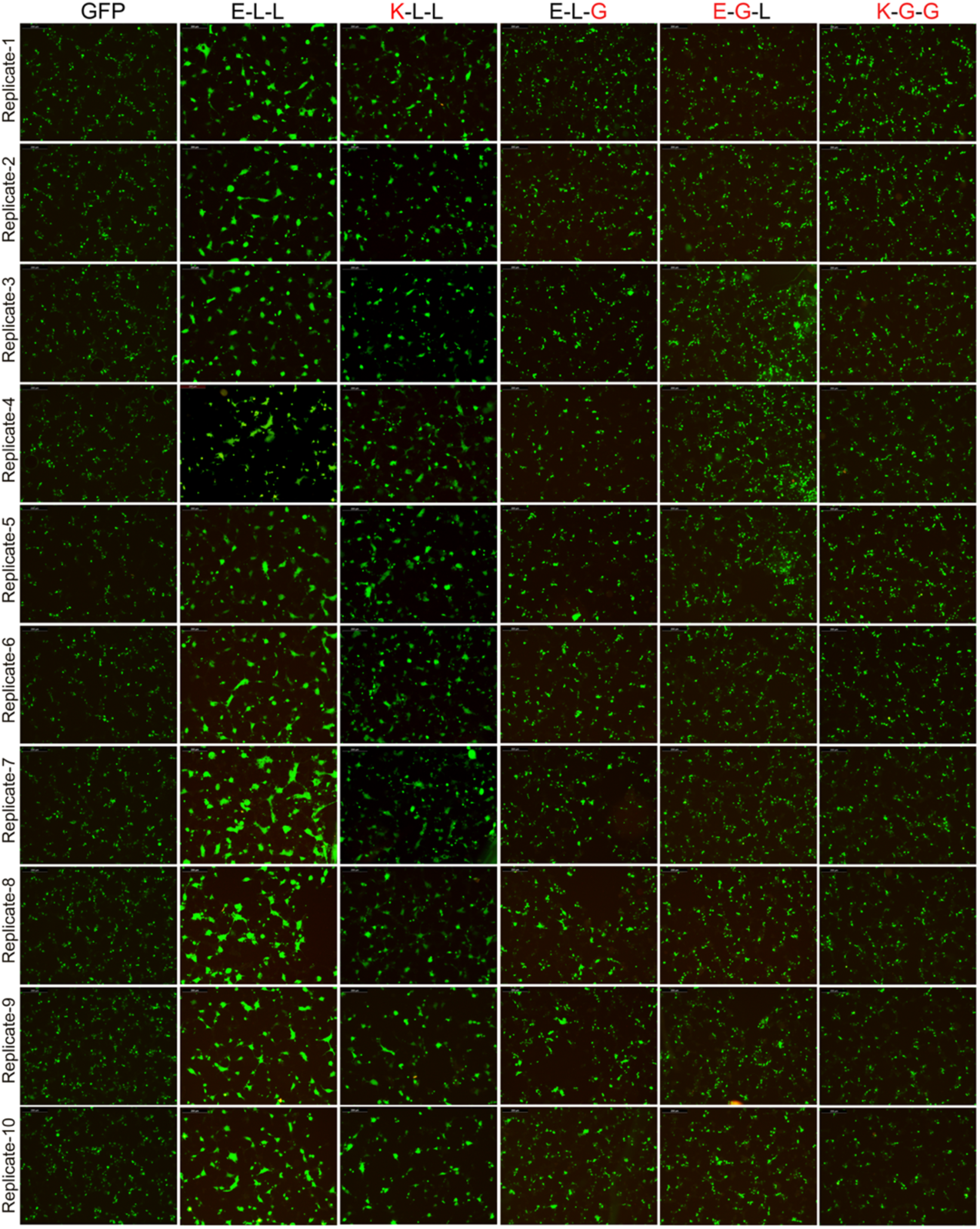
WT or mutant spike proteins were tested for their ability to trigger syncytium formation in cells overexpressing the ACE2 receptor. HEK293T cells overexpressing SARS-CoV-2 spike (WT and mutants) and eGFP proteins were co-cultured with HEK293T cells overexpressing ACE2 receptors for twelve hours before they were fixed and imaged using a fluorescence microscope. Ten different filed were imaged and are of GFP positive cells were quantified using Image J software (quantification shown in figure 2M).

**Figure S4.**
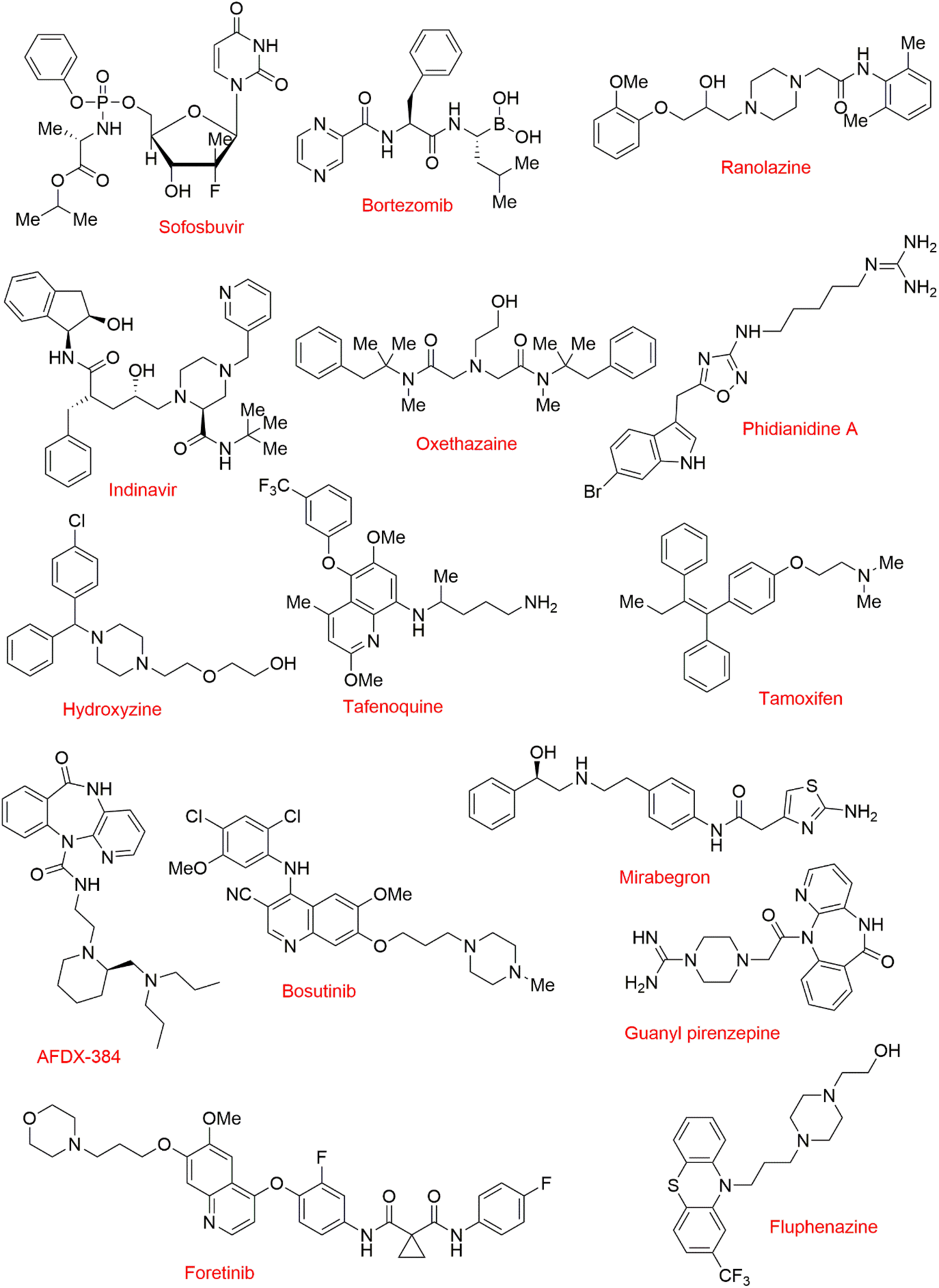
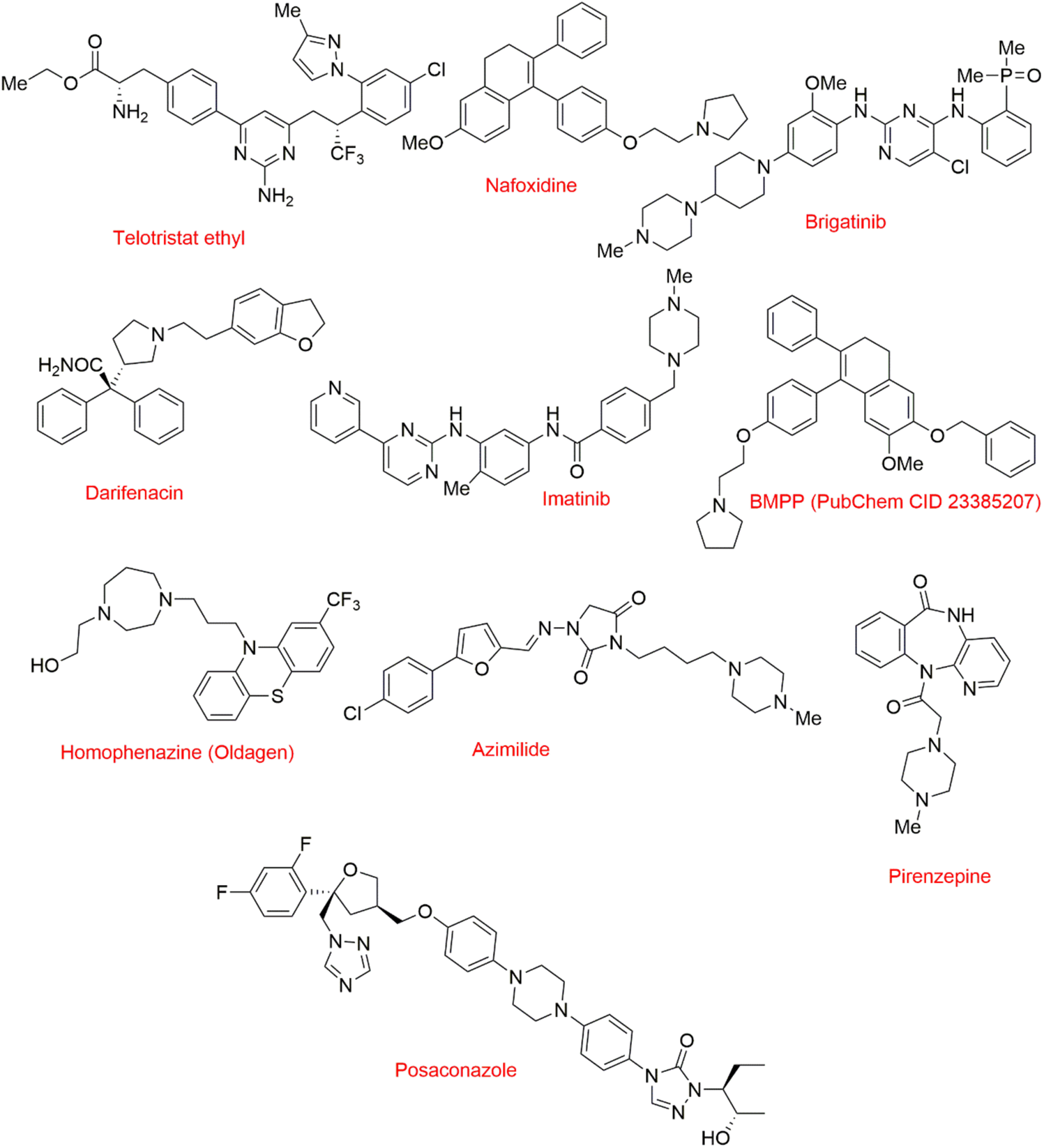
Chemical structures of the 25 selected drug candidates. Images were created using Chemdraw software.

**Figure S5.**
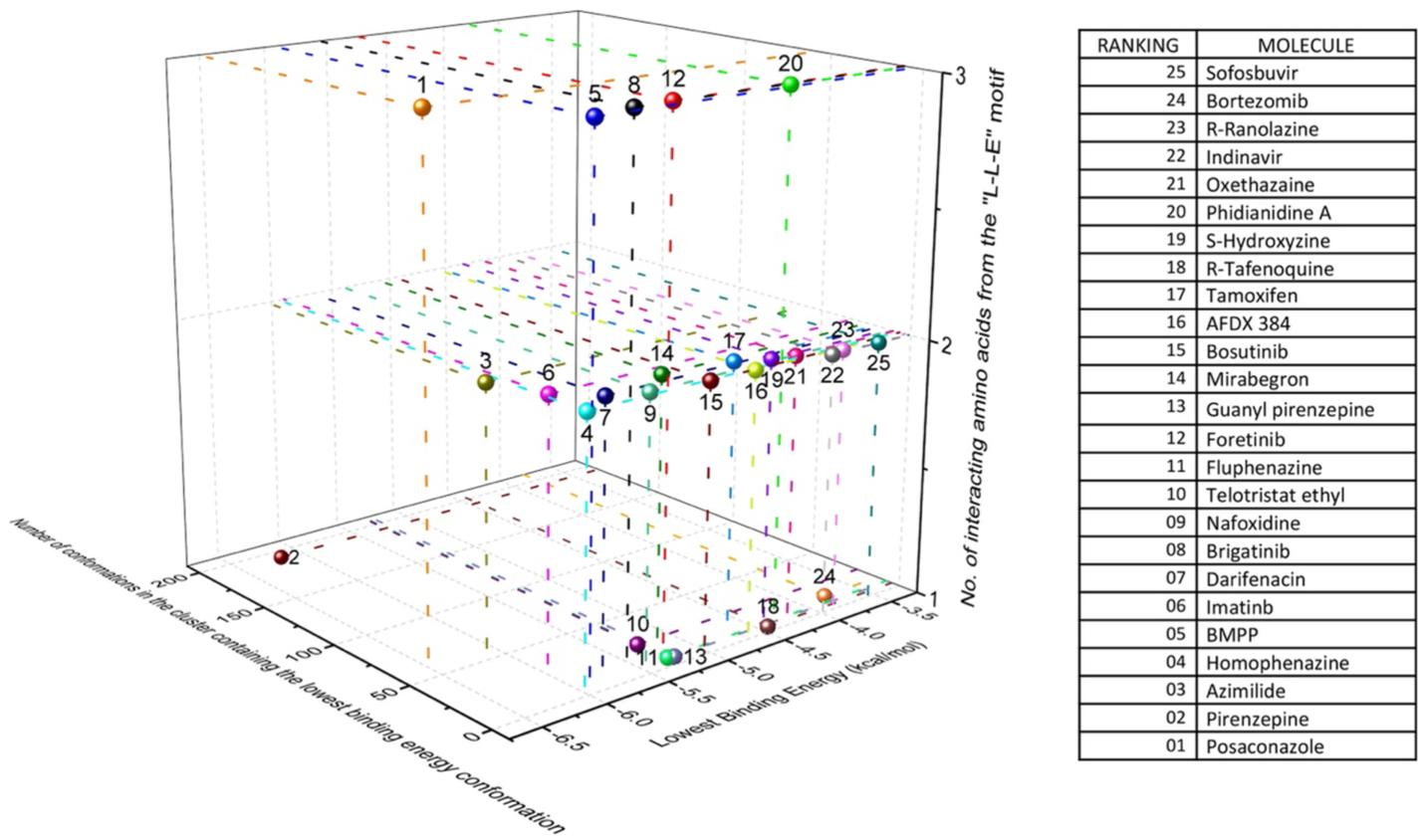
Three-dimensional representation of comparative assessment drug molecules based upon three parameters. (i) Their lowest binding energies towards the target pocket present in the protein (ii) number of conformations present in the cluster containing the lowest binding energy conformation and (iii) number of amino acids amongst the E-L-L motif with which the lowest binding energy conformation interacts.

**Figure S6.**
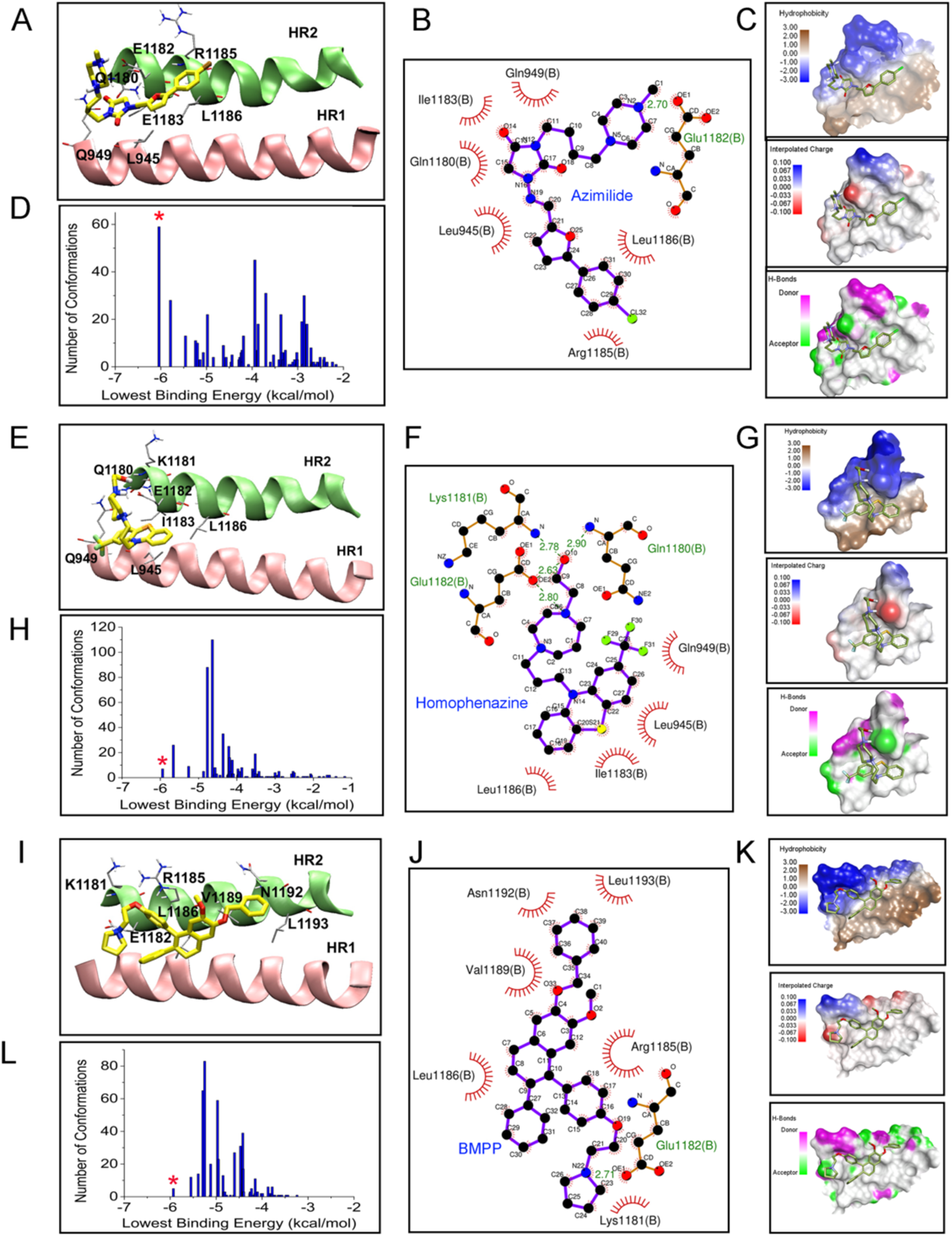
Images A-D corresponds to the lowest binding energy conformation of azimilide; E-H corresponds to the lowest binding energy conformation of homophenazine and I-L corresponds to the lowest binding energy conformation of BMPP. A, E, and I represent the binding pocket for each of the corresponding ligand conformations with HR1 and HR2 helix shown as cartoon image, ligand molecule, and the amino acid side chains shown in stick model. In images, C, G, and K, hydrophobicity, hydrogen bond donor/acceptor and interpolated charges on the protein surface for each corresponding ligand conformation have been mapped. In D, H, and L show cluster analysis for each of the ligands where the asterisk marks the particular cluster to which the corresponding ligand conformation belongs; B, F, and J are the ligplot images showing the corresponding ligand-protein interaction in two-dimension.

**Figure S7.**
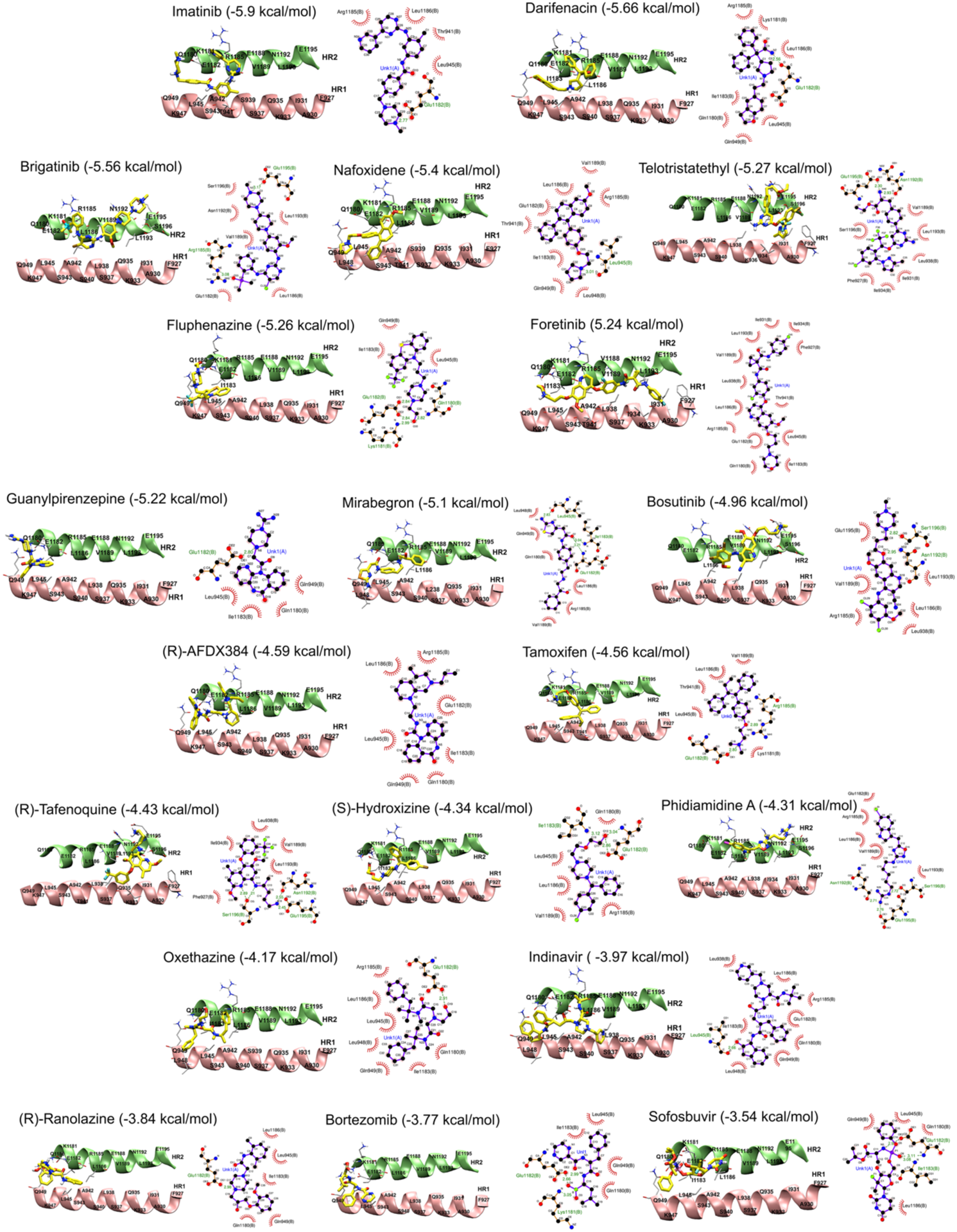
Ligand binding pocket in the protein with cartoon representation of HR1 and HR2 helices and corresponding ligplots showing protein-ligand interactions are presented for 20 selected drug molecules.

**Figure S8.**
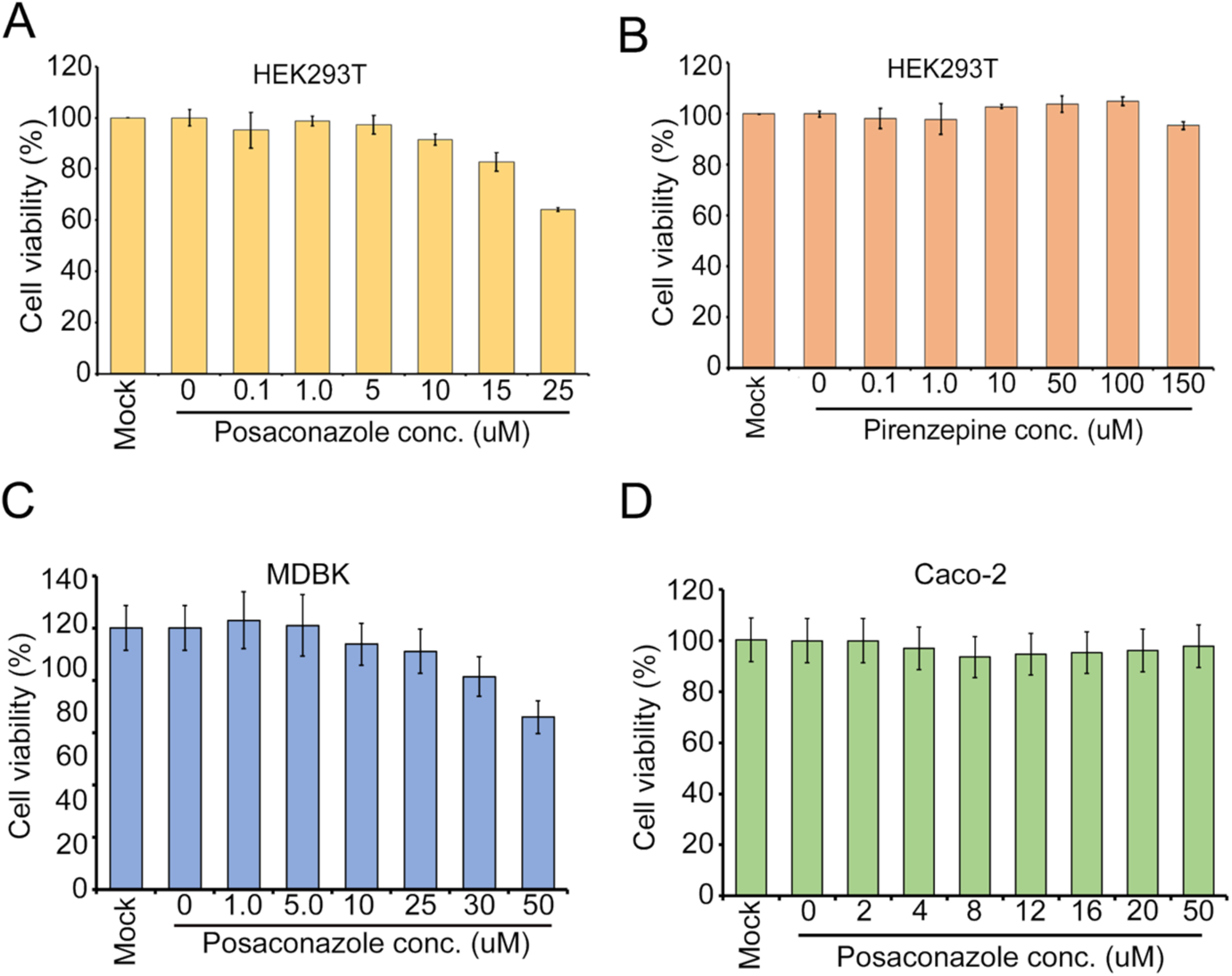
Cytotoxicity of drugs. (A-B) HEK293T cells or (C) MDCK cells or (D) Caco-2 cells were treated with different concentrations of Posaconazole or Pirenzepine for 24h as indicated. Cytotoxicity was determined by MTT assay.

**Figure S9.**
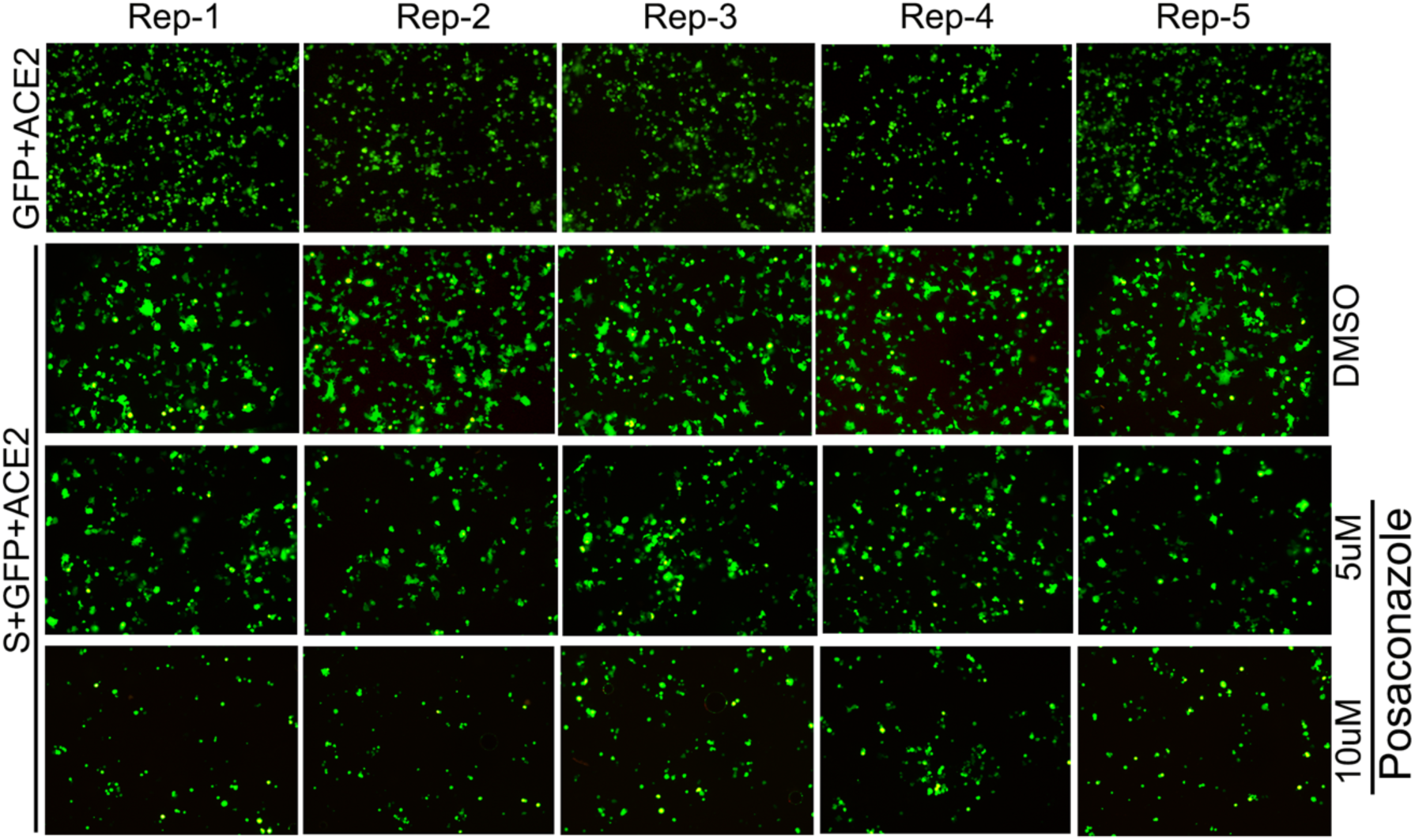
Cells expressing SARS-CoV-2 spike and GFP proteins were treated with different concentrations of posaconazole or with the solvent before and during their coculturing with HEK293T-ACE2 cells for 12 hours and syncytium formation was monitored under a fluorescence microscope.

**Figure S10.**
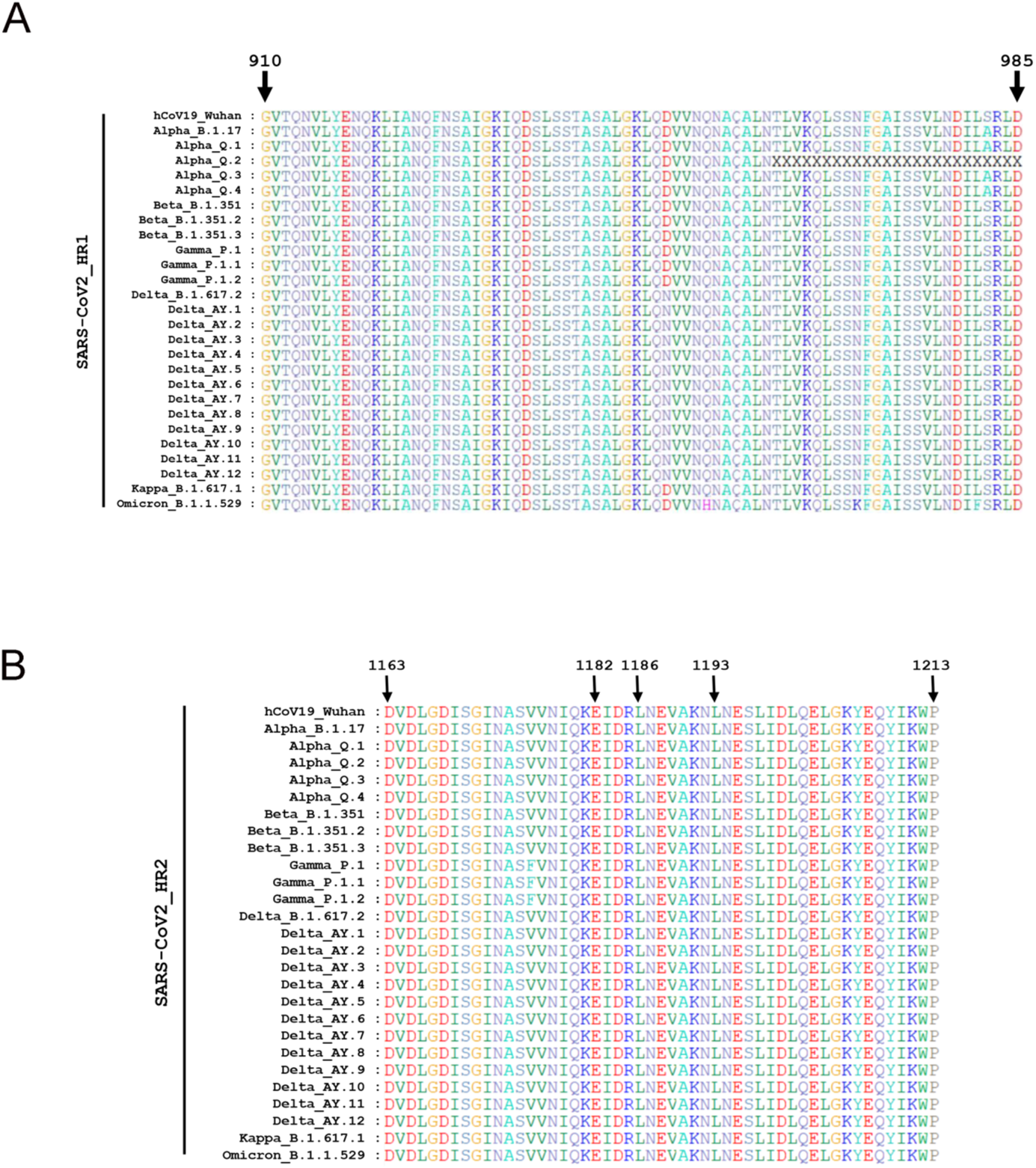
Multiple sequence alignment of spike HR1 (A) and HR2 (B) domains of S protein from different variants & sub-variants of SARS-CoV-2, highlighting overall conservation of the amino acid sequences. The sequences were retrieved from the GISAID database and aligned using Bioedit software.

**Figure S11.**
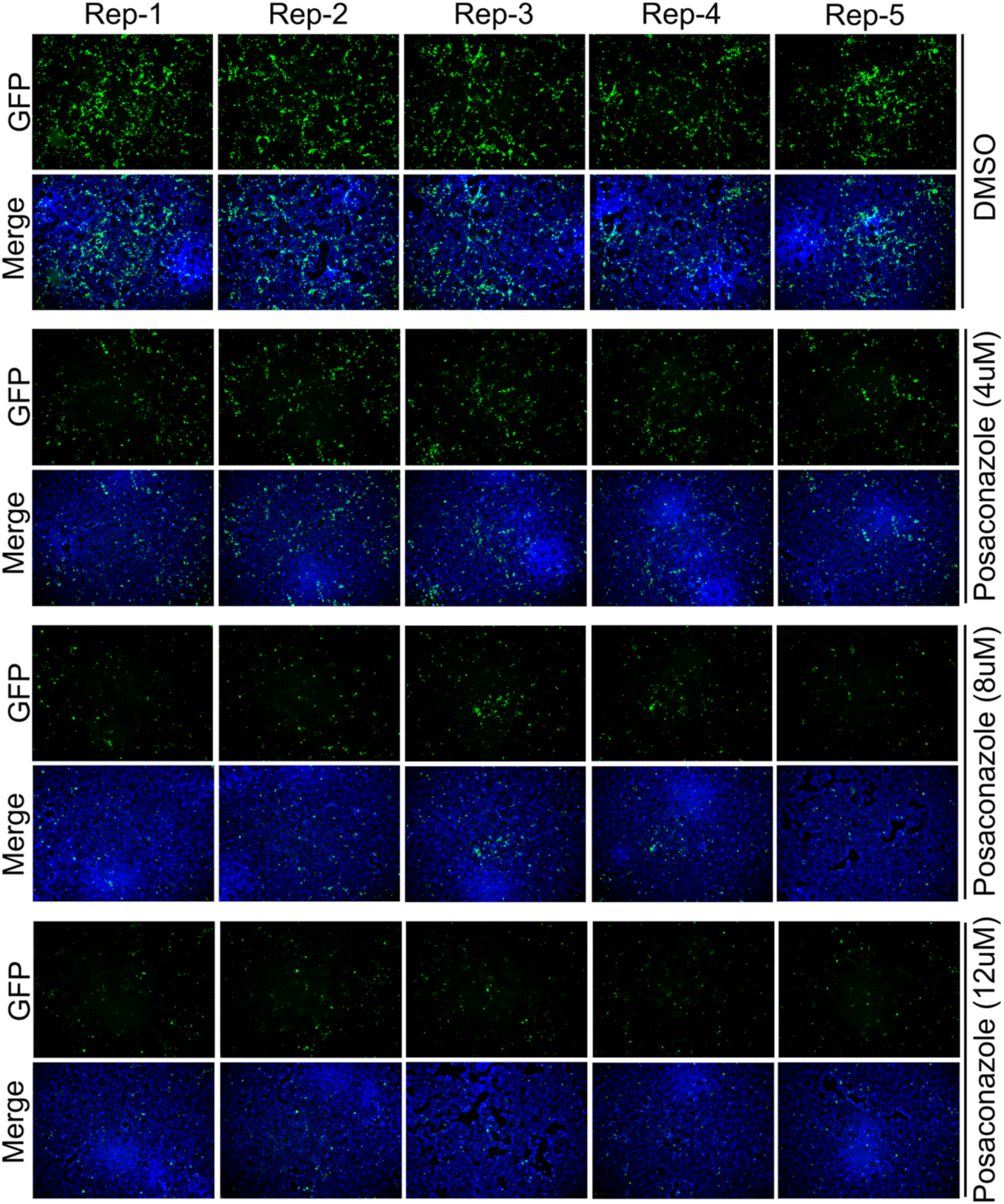
(A) Caco-2 cells pretreated for 12 h with DMSO or the indicated concentrations of posaconazole were infected with SARS-CoV-2-mNG virus (multiplicity of infection, MOI of 0.1) pretreated for 1 h with the corresponding concentrations of the drug. 48 h after infection, the mNG and DAPI signals were visualized from five different fields of each set and quantitated using ImageJ software (quantification shown in Figure 5B).

**Figure S12.**
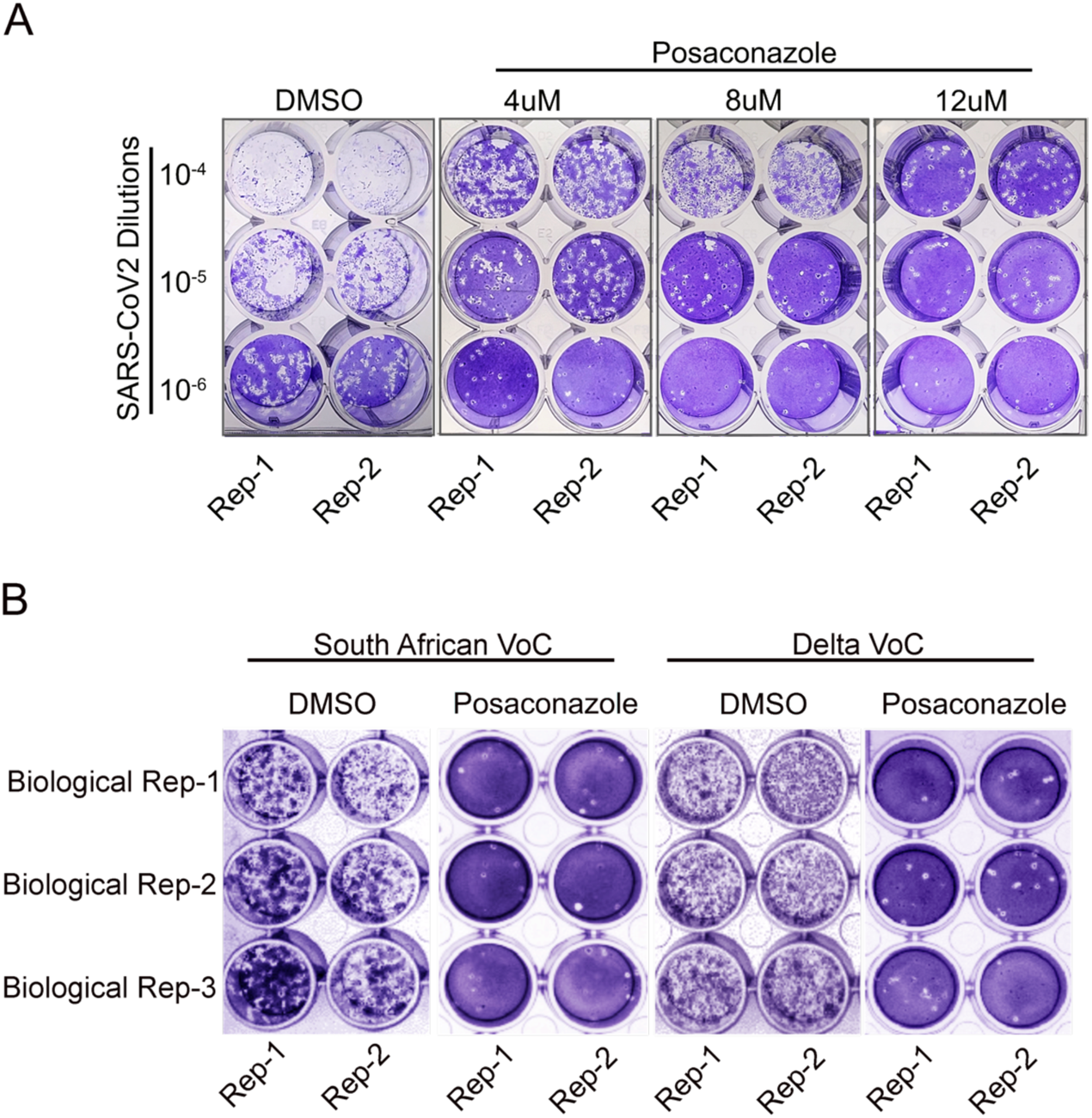
Posaconazole treatment reduces the titer of WT and different variants of SARS-CoV-2. (A) Caco-2 cells pretreated for 1 h with DMSO or the indicated concentrations of posaconazole were infected with WT SARS-CoV-2 (2019-nCoV/USA_WA1/2020) virus (multiplicity of infection, MOI of 0.1) pretreated for 1 h with the corresponding concentrations of the drug. 48 h after infection supernatants were harvested and used for plaque assay in Vero E6-ACE2-TMPRSS2 cells. (B) Indicated VoCs of SARS-CoV-2 and Caco-2 cells were pretreated with vehicle control or with 8uM of Posaconazole before infecting the cells with 10^-6^ dilutions of the viruses. Supernatants were harvested 48 hours after infection and titrated using plaque assay.

**Figure S13.**
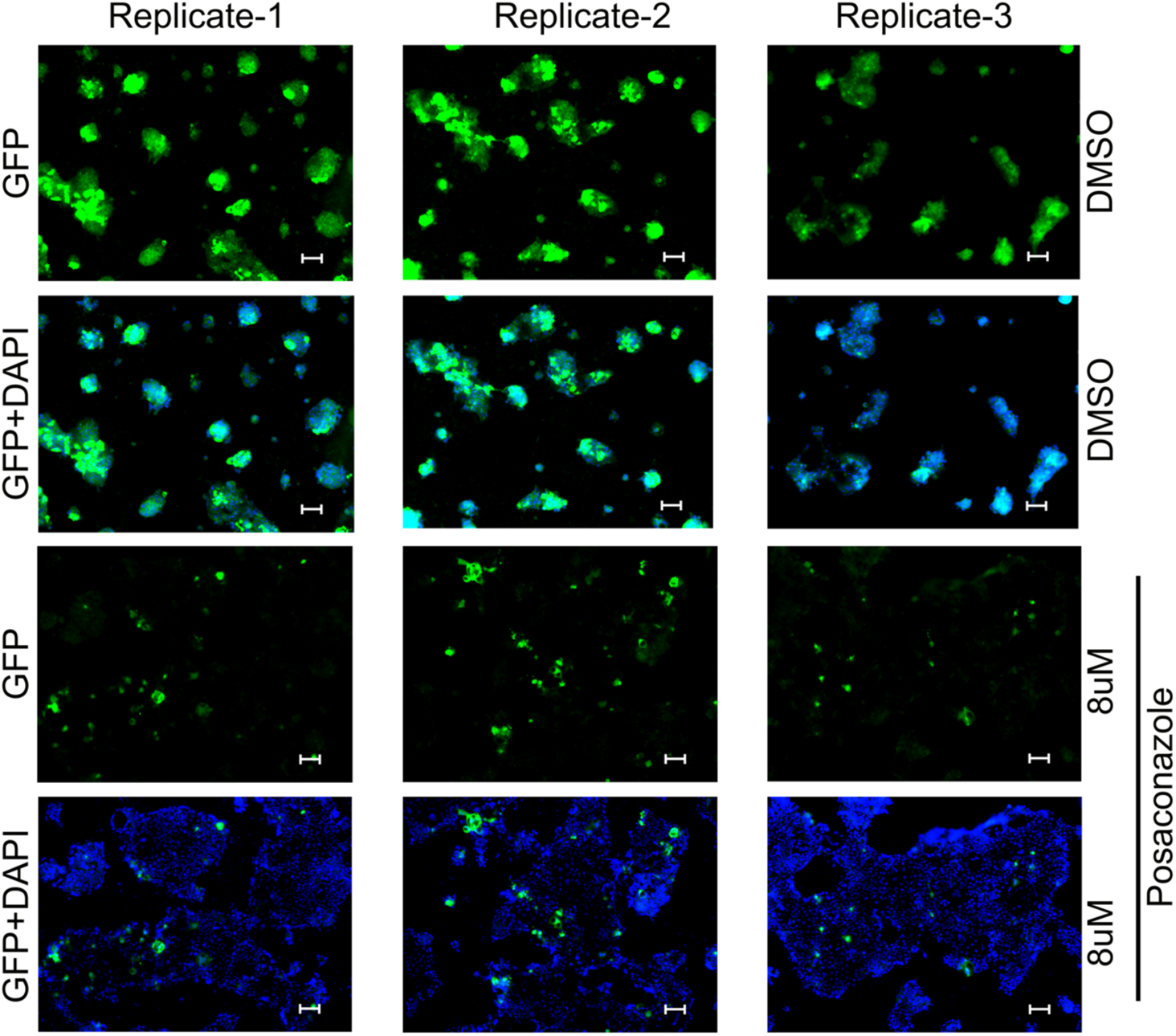
Caco-2 cells pretreated with posaconazole (8 µM) were infected with pretreated SARS-CoV-2-mNG (MOI 0.1) and visualized at 40X for syncytia formation. Three different fields were imaged for each set to measure the area of nNG positive cells using ImageJ software.

**Figure S14.**
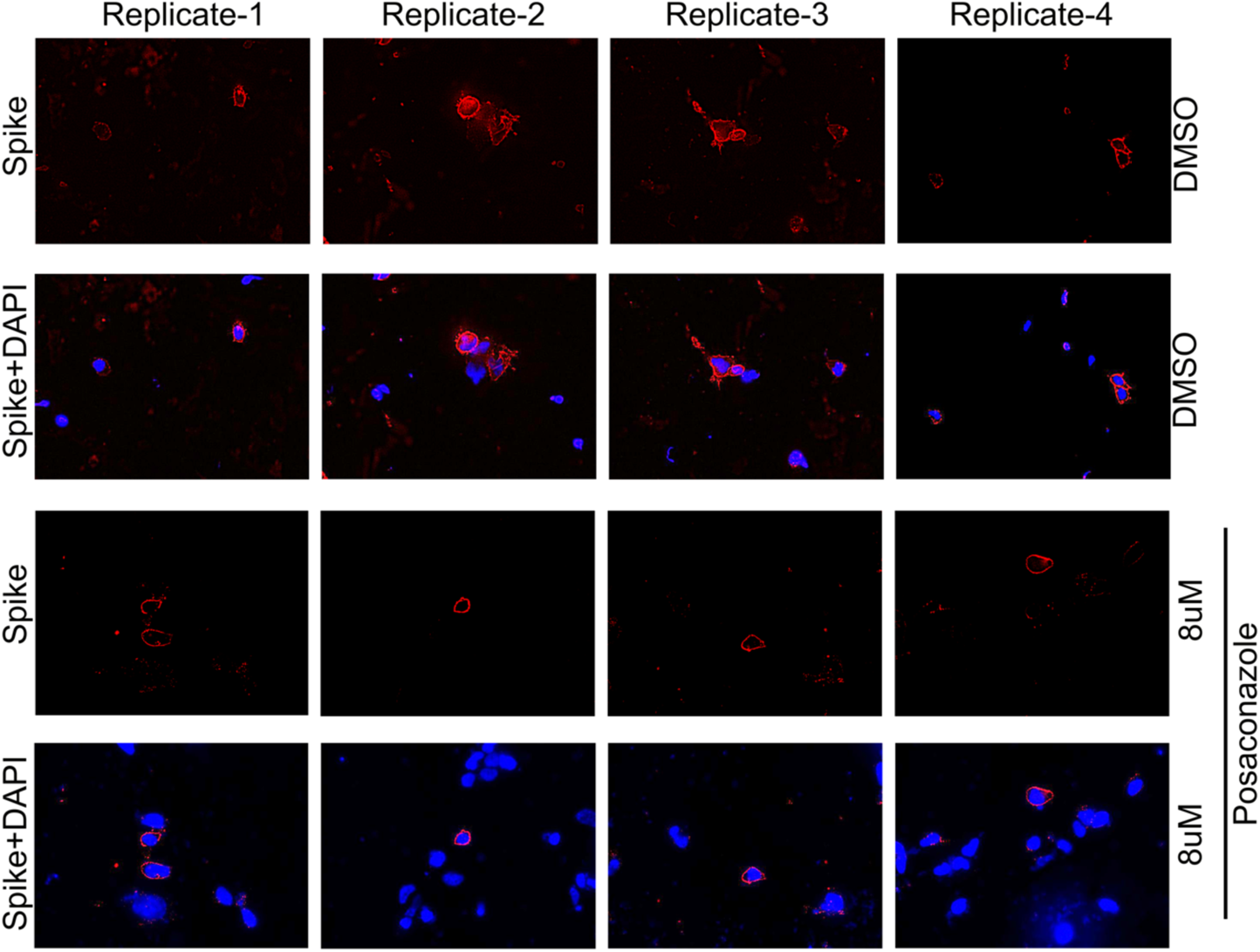
Surface immunofluorescence assay was performed on Caco-2 cells infected with WT SARS-CoV-2 in presence of DMSO (mock) or posaconazole (8 µM). Four different fields were visualized for each set.

**Figure S15.**
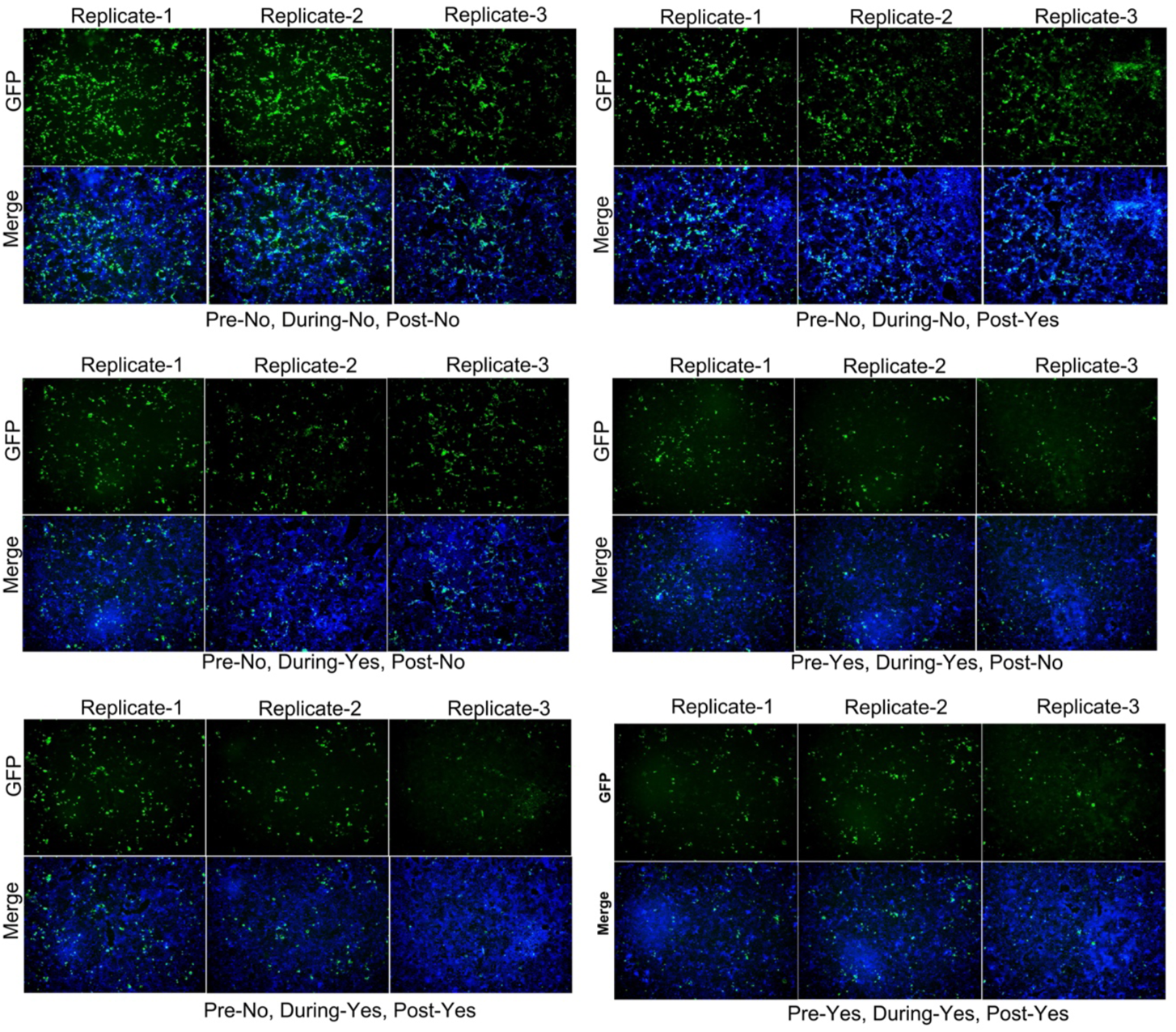
Caco-2 cells and SARS-CoV-2-mNG (MOI 0.1) were treated with Posaconazole (8uM) at indicated times while performing the infection. mNG and DAPI signals were visualized from three independent filed for each set and used for quantitative analysis using ImageJ software.

**Table S1.**
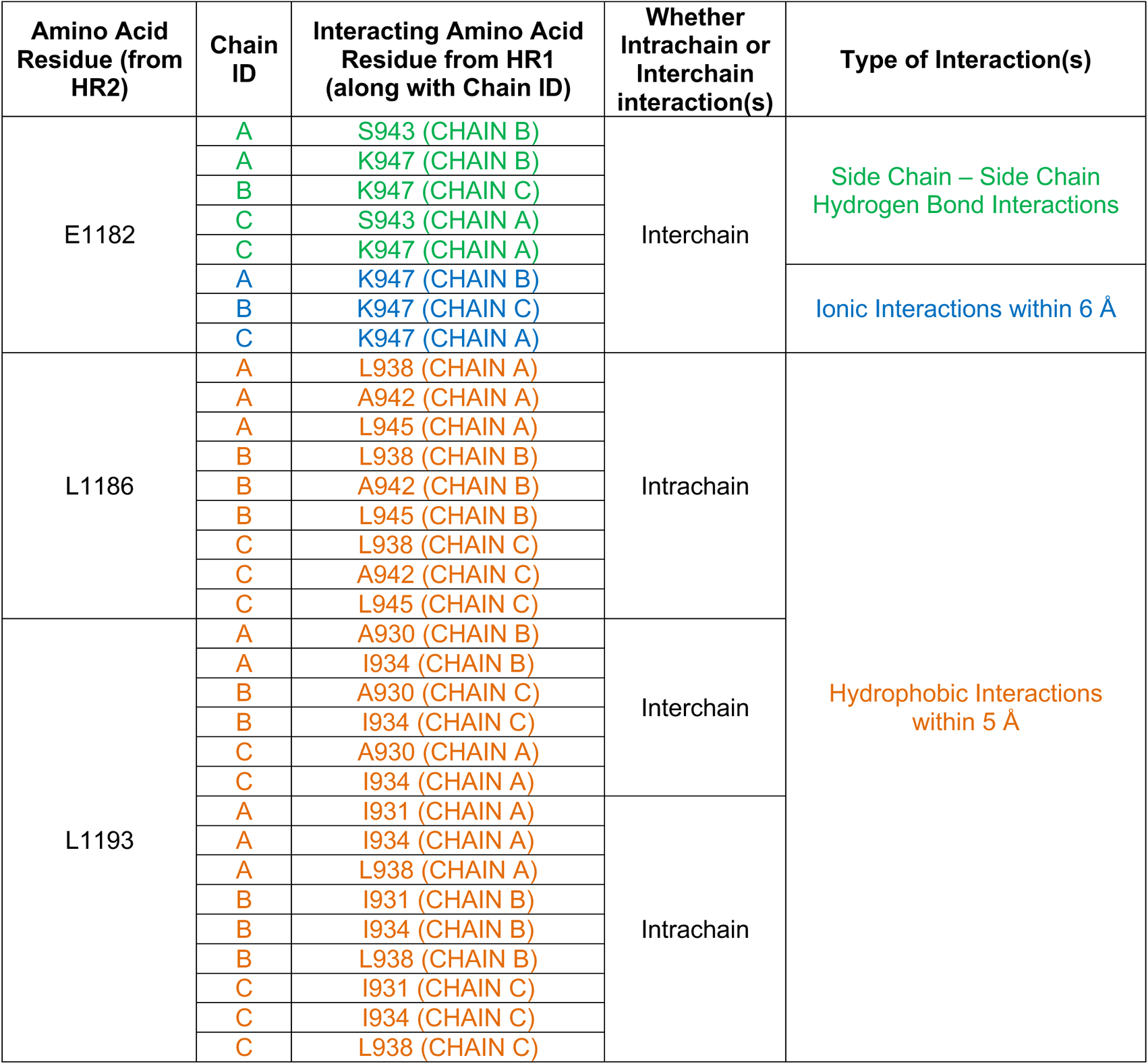
Different intra and intermolecular interactions between HR1 and HR2 domains within the 6-HB structure involving the E1182, L1186, and L1193 amino acid residues of the HR2. The stature (PDB ID:6LXT) was analyzed using the PIC web server.

**Table S2.** Twenty drug candidates (excluding top 5) and their interaction profiles with the HR1-HR2 bis-peptide in their lowest binding energy conformation.

| MOLECULES | CLINICAL USE | LOWEST BINDING ENERGY (kcal/mol) | INTERACTING AMINO ACID RESIDUES |  | NUMBER OF CONFORMATIONS IN THE CLUSTER CONTAINING THE LOWEST BINDING ENERGY CONFORMATION. |
| --- | --- | --- | --- | --- | --- |
|  |  |  | Interacting through H-bonding and/or ionic interactions (for charged moieties) | Interacting through hydrophobic interactions |  |
| **Sofosbuvir**<br>FDA approved | Hepatitis C medication | -3.54 | **E 1182**, I 1183, | Q 949, L 945, Q 1180, **L 1186** | 1 |
| **Bortezomib**<br>FDA approved | Anti-cancer medication | -3.77 | **E 1182**, K 1181 | Q 949, L 945, Q 1180, 1183 | 12 |
| **Ranolazine**<br>FDA approved<br>(racemic drug)<br>R-configuration was used for docking | Heart related chest pain medication | -3.84 | **E 1182** | **L 1186**, L 945, I 1183, Q 949, Q 1180 | 3 |
| **Indinavir**<br>FDA approved | Anti-retroviral medication for HIV-AIDS | -3.97 | L 945 | L 938, **L 1186**, R 1185, **E 1182**, Q 1180, Q 949, L 948, I 1183 | 1 |
| **Oxethazaine**<br>FDA approved | Peptic ulcer disease or esophagitis medication | -4.17 | **E 1182** | R 1185, Q 1180, I 1183, Q 949, L 948, L 945, **L 1186** | 9 |
| **Phidianidine A**<br>(Marine natural product) | Anti-foulants | -4.31 | N 1192, S 1196, E 1195 | **E 1182**, **L 1193**, V 1189, **L 1186**, R 1185 | 8 |
| **Hydroxyzine**<br>FDA approved<br>(racemic drug)<br>S-configuration was used for docking) | Antihistamine used for allergy medication | -4.34 | **E 1182**, I 1183 | Q 1180, R 1185, V 1189, **L 1186**, L 945 | 12 |
| **Tafenoquine**<br>FDA approved<br>(racemic drug)<br>R-configuration was used for docking) | Medication used to prevent and to treat malaria | -4.43 | N 1192, S 1196, E 1195 | L 938, V 1189, **L 1193**, F 927, I 934 | 4 |
| **Tamoxifen**<br>FDA approved | Selective estrogen receptor modulator used for medication to prevent breast cancer | -4.57 | **E 1182** | **L 1186**, R 1185, L 938, V 1189, K 1181 | 19 |
| **AFDX 384**<br>Under trial | Selective antagonist of the muscarinic acetylcholine receptors | -4.59 | - | R 1185, **E 1182**, I 1183, Q 1180, Q 949, L 945, **L 1186** | 5 |
| **Bosutinib**<br>FDA approved | BCR-ABL and src tyrosine kinase inhibitor used for the medication of chronic myelogenous leukemia | -4.96 | S 1196, N 1192 | **L 1193**, **L 1186**, L 938, R 1185, V 1189, E 1195 | 6 |
| **Mirabegron**<br>FDA approved | Medication used to treat overactive bladder | -5.10 | L 945, I 1183, **E 1182** | L 948, **L 1186**, R 1185, V 1189, Q 1180, E 949 | 24 |
| **Guanyl pirenzepine** | Muscarinic antagonist | -5.22 | **E 1182** | L 945, I 1183, Q 1180, Q 949 | 6 |
| **Foretinib**<br>Under trial | Drug candidate for the treatment of cancer | -5.24 | - | I 931, I 934, F 927, T 941, L 945, I 1183, Q 1180, **E 1182**, R 1185, **L 1186**, L 938, V 1189, **L 1193** | 11 |
| **Fluphenazine**<br>FDA approved | Antipsychotic medication | -5.26 | Q 1180, K 1181, **E 1182** | Q 949, L 945, I 1183 | 7 |
| **Telotristat ethyl**<br>FDA approved | Medication for adults with diarrhea associated with carcinoid syndrome | -5.27 | E 1195, N 1192 | V 1189, **L 1193**, L 938, I 931, I 934, F 927, S 1196, | 24 |
| **Nafoxidine**<br>Under trial | Nonsteroidal selective estrogen receptor modulator | -5.40 | L 945 | V 1189, R 1185, L 948, Q 949, I 1183, T 941, **E 1182**, **L 1186** | 10 |
| **Brigatinib**<br>FDA approved | Anaplastic lymphoma kinase (ALK) and epidermal growth factor receptor (EGFR) inhibitor used in the medication for cancer | -5.56 | R 1185, E 1195 | **L 1193**, **L 1186**, **E 1182**, V 1189, N 1192, S 1196 | 10 |
| **Darifenacin**<br>FDA approved | M<sub>3</sub> muscarinic acetylcholine receptor blocker used for the medication of overactive bladder | -5.66 | **E 1182** | R 1185, K 1181, **L 1186**, L 945, Q 949, Q 1180, I 1183 | 17 |
| **Imatinib**<br>FDA approved | Tyrosine Kinase inhibitor used for the medication of cancer | -5.90 | **E 1182** | R 1185, **L 1186**, T 941, L 945 | 32 |

**Table S3.** Hydrogen Bonding of Posaconazole in its lowest binding conformation

| Serial Number | Moiety from Posaconazole | Moiety from HR2 | Distance (Donor...Acceptor) | Angle (<Donor-H-Acceptor) |
| --- | --- | --- | --- | --- |
| 1. | OH (O1) (Donor) | Carboxylate oxygen (OE2) of E1182 (Acceptor) | 2.75Å | 143.5° |
| 2. | OH (O1) (Donor) | Backbone N of E1182 (Acceptor) | 2.97Å | 128.1° |
| 3. | O1 (Acceptor) | Backbone NH of I1183 (Donor) | 3.20Å | 150.4° |
| 4. | Triazole nitrogen (N7) (Acceptor) | –OH (OG) of S1196 (Donor) | 3.02Å | 153.2° |

**Table S4.**
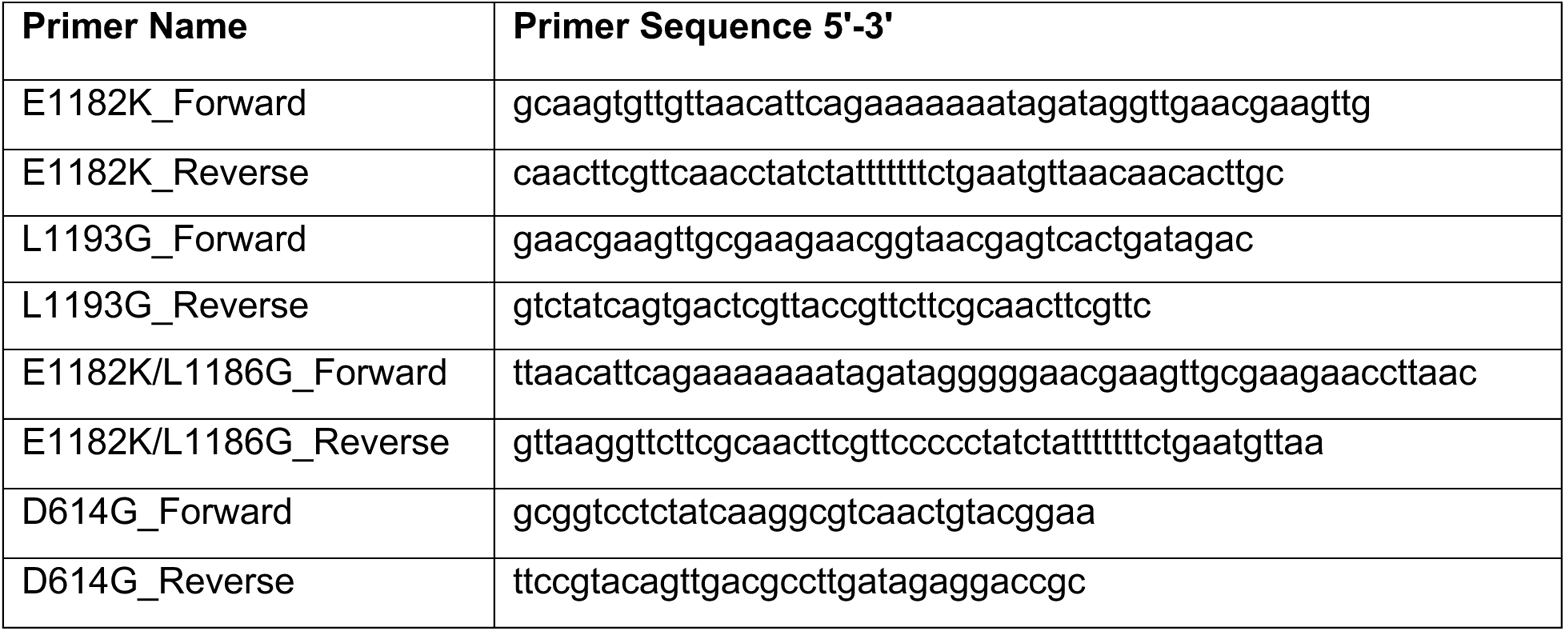
Primer list for site-directed mutagenesis in the SARS-CoV-2 spike ORF.

